# Genome evolution and transcriptome plasticity associated with adaptation to monocot and eudicot plants in *Colletotrichum* fungi

**DOI:** 10.1101/2022.09.22.508453

**Authors:** Riccardo Baroncelli, José F. Cobo-Díaz, Tiziano Benocci, Mao Peng, Evy Battaglia, Sajeet Haridas, William Andreopoulos, Kurt LaButti, Jasmyn Pangilinan, Anna Lipzen, Maxim Koriabine, Diane Bauer, Gaetan Le Floch, Miia R. Mäkelä, Elodie Drula, Bernard Henrissat, Igor V. Grigoriev, Jo Anne Crouch, Ronald P. de Vries, Serenella A. Sukno, Michael R. Thon

## Abstract

*Colletotrichum* fungi infect a wide diversity of monocot and eudicot hosts, causing plant diseases on almost all economically important crops worldwide. In addition to its economic impact, *Colletotrichum* is a suitable model for the study of gene family evolution on a fine scale to uncover events in the genome that are associated with the evolution of biological characters important for host interactions. Here we present the genome sequences of 30 *Colletotrichum* species, 18 of them newly sequenced, covering the taxonomic diversity within the genus. A time-calibrated tree revealed that the *Colletotrichum* ancestor diverged in the late Cretaceous around 70 million years ago (mya) in parallel with the diversification of flowering plants. We provide evidence of independent host jumps from eudicots to monocots during the evolution of this pathogen, coinciding with a progressive shrinking of the degradative arsenal and expansions in lineage specific genes. Comparative transcriptomics of four reference species with different evolutionary histories and adapted to different hosts revealed similarity in gene content but differences in the modulation of their transcription profiles. Only a few orthologs show similar expression profiles on different plant cell walls. Combining genome sequences and expression profiles we identified a set of core genes, such as specific transcription factors, involved in plant cell wall degradation in *Colletotrichum*.Together, these results indicate that the ancestral *Colletotrichum* were associated with eudicot plants and certain branches progressively adapted to different monocot hosts, reshaping part of the degradative and transcriptional arsenal.

## INTRODUCTION

The plant cell wall (PCW) consists of many different polysaccharides that are attached not only to each other through a variety of linkages, but also to the aromatic polymer lignin, providing the main strength and structure for the PCW. In addition, PCWs are determinants of immune responses since modification of their composition affect disease resistance and fitness on plants^1–3^.

Fungi have developed a complex and efficient plant biomass degrading machinery, involved not only in saprotrophic degradation, but also in other types of plant-fungal interactions, such as pathogenicity, symbiosis, and parasitism. A common characteristic of plant pathogenic fungi is the need to cross the PCW, an important barrier against pathogen invasion. In this context, the PCW can be seen as one of the first levels of the arms race between the pathogen and the host, but also as a complex ecological niche where the fungi retrieve most of the nutrients from the host during infection. To release the monomers present in these complex plant structures, fungi need to simultaneously secrete several plant biomass degrading enzymes, consisting of a large set of hydrolytic and oxidative enzymes^2^. Plants protect themselves against destruction of their cell walls by producing proteins that inhibit microbial cell wall degrading enzymes (CWDEs), e.g., inhibitors of pectin-degrading enzymes are common in eudicots and non-commelinoid monocots, and inhibitors of xylan-degrading enzymes are common in grasses^4^. The production of these inhibitors by plants has, in turn, driven the evolution of some CWDE groups of phytopathogenic fungi toward inhibitor-resistant enzymes^5^. In some phytopathogenic fungi, there is evidence for production of different amounts of specific CWDEs, depending on whether the plant host is a monocot or eudicot^6–8^.

*Colletotrichum* spp. are among the most scientifically and economically important plant pathogenic fungi^9^. Outbreaks of *Colletotrichum* spp. have been devastating^10^ and, besides the economic impact, several *Colletotrichum* spp. have been widely used as model systems for studies of plant pathology and fungal–plant interactions^11, 12^. *Colletotrichum* species have an endophytic and/or a pathogenic association with over 3,200 species of monocot and eudicot plants^13^. Many species within this genus have a hemibiotrophic infection strategy, with an initial biotrophic phase followed by a necrotrophic phase characterized by fast-growing hyphae that destroy host tissues^6^. There is a high variety of host-specific relationships among *Colletotrichum* species. Some species show a one-to-one relationship with a specific host while other species infect a wide range of hosts^6, 12, 14–16^. The biological diversity of *Colletotrichum* and the presence of very closely related species with different host range makes this genus a suitable model to investigate genomic signatures associated with the evolution of phenotypic characters important for host interactions such as those involved in PCW degradation.

Since the first genome sequences of fungi became available, researchers have been analyzing gene content and genomic features to find associations that may explain the differences in fungal lifestyles and varying patterns are beginning to emerge^6, 17^. A unique genome provides information on gene content, encoded proteins and genome structure, but comparative studies between closely related species with different biological features are necessary to elucidate which genetic elements could be involved in a specific trait and to study their evolution^18, 19^. The loss of lineage specific effector protein candidates (LSECs) was also detected by comparative genomic studies in powdery mildew fungi^20^. In contrast, gene loss or gain in families such as those encoding CAZymes and proteases could be associated with host range in *Colletotrichum* species^21^. The similar repertoires of CAZymes and secreted proteases found in relatively distant members of the *C. acutatum* and *C. gloeosporioides* species complexes suggest a recent and independent acquisition of this enzymatic arsenal or a progressive lost during the host specialization process^6, 21, 22^. While genome studies are useful tools to identify putative genes and to perform evolutionary analyses, transcriptomic data is required to better understand the genes involved in a complex process such as PCW interaction.

Plant pathogenic fungi have a close interaction with the PCW and plants have evolved to recognize external attacks through the degradation of the PCW itself. This is especially true for hemibiotrophic plant pathogens as they interact with the PCW twice: initially when they enter the cell and later when they gain nutrients from it. This complexity is reflected by the wide arsenal of CAZymes encoded by *Colletotrichum* spp. being one of the most diverse in the fungal kingdom. In this work, we used a comparative genomics and transcriptomics approach to identify genes involved in the interaction between *Colletotrichum* spp. and PCW, and evolutionary analyses to gain a better understanding of adaptation and specialization of these fungi to different PCWs. Phylogenetic analyses revealed that the ancestral *Colletotrichum* was associated with eudicots and that at least 3 independent jumps to monocots occurred. We also found that monocot associated *Colletotrichum* species have undergone specific gene losses in PCW degrading enzyme families and expansions in lineage specific genes.

Comparing four different species we also found that, despite maintaining a similar gene content, *Colletotrichum* species show strong differences in gene modulation, possibly due to adaptation to different host substrates.

## RESULTS

### The common ancestor of *Colletotrichum* parasitized eudicots and specific lineages jumped independently to monocots

In this study, we present a comparative genomic analysis of 30 species from the genus *Colletotrichum*. Eleven of these (*C. cereale*, *C. eremochloae*, *C. sublineola*, *C. graminicola*, *C. falcatum*, *C. navitas*, *C. caudatum*, *C. somersetensis*, *C. zoysiae*, *C. orchidophilum* and *C. phormii*) are pathogens specialized to different taxonomic groups of monocots; seventeen (*C. orbiculare*, *C. nupharicola*, *C. higginsianum*, *C. tofieldiae*, *C. salicis*, *C. godetiae*, *C. acutatum sensu stricto*, *C. fioriniae*, *C. abscissum*, *C. lupini*, *C. tamarilloi*, *C. costaricense*, *C. cuscutae*, *C. paranaense*, *C. melonis, C. nymphaeae* and *C. simmondsii*) have been associated only with eudicots while two of them (*C. chlorophyti* and *C. incanum*) are capable of infecting plants that belong to both groups.

The analyzed genomes showed a large variation in size, ranging from 44.20 Mb in *C. caudatum* to 89.65 in *C. orbiculare* (Figure 1). While a large variation at the genus level was already reported^23^ (more than 50% in our dataset) these results highlight an unexpected variation of more than 30 Mb (39%) between closely related species such as *C. cuscutae* and *C. paranaense*. These two species belong to the Acutatum species complex and have been recognized as separate taxa only recently. As a general trend, species with bigger genomes are characterized by a lower GC content. This evidence coupled with the lack of correlation between genome size and number of predicted genes suggest that the genome size in *Colletotrichum* is mainly correlated with the amount of non-coding DNA (Figures 1 and 2). Phylogenomic analyses calibrated with three fungal fossils show age estimates for *Colletotrichum* spp. and enable the identification of time frames of specific evolutionary events (Figure 1).

**Figure 1.**
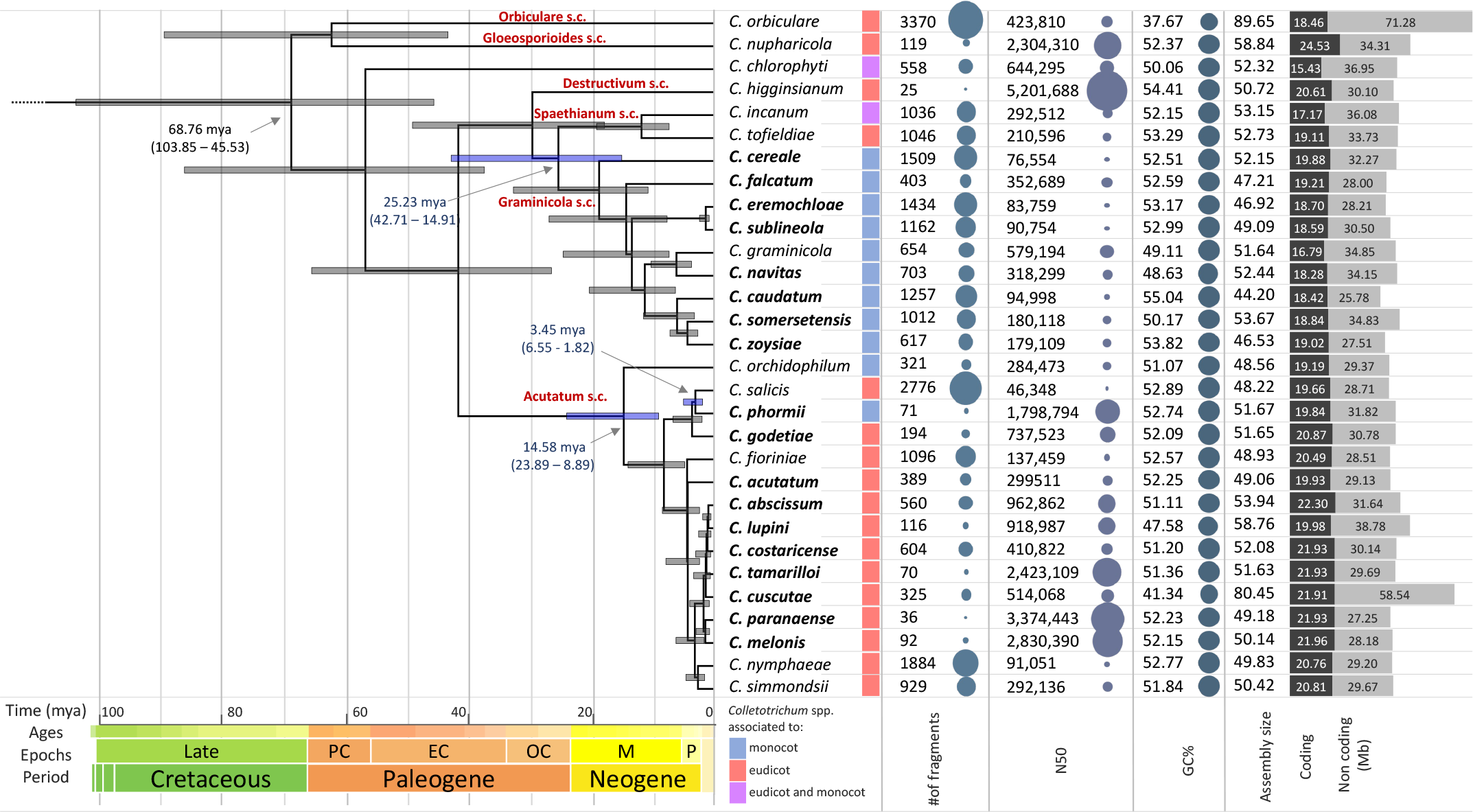
A timetree inferred by the RelTime method to the *Colletotrichum* phylogenomic tree. The branch lengths were calculated using the Ordinary Least Squares method. All nodes are supported by Bayesian posterior probability of 1.00. Bars around each node represent 95% confidence intervals and light blue bars represent the three host jumps from eudicot to monocot. This analysis involved 127 amino acid sequences and a total of 124023 sites. *Colletotrichum* species complexes are indicated in red. Genomes sequenced in the present study are highlighted in bold. On the right side four bubble plots illustrating assembly size, GC content and assembly fragmentation parameters (number of contigs and N50 value) and are reported in the right side. The bubble sizes have been scaled to each panel and are not comparable across panels. Gray bar diagram on the right reports the size of coding and non-coding regions.

**Figure 2.**
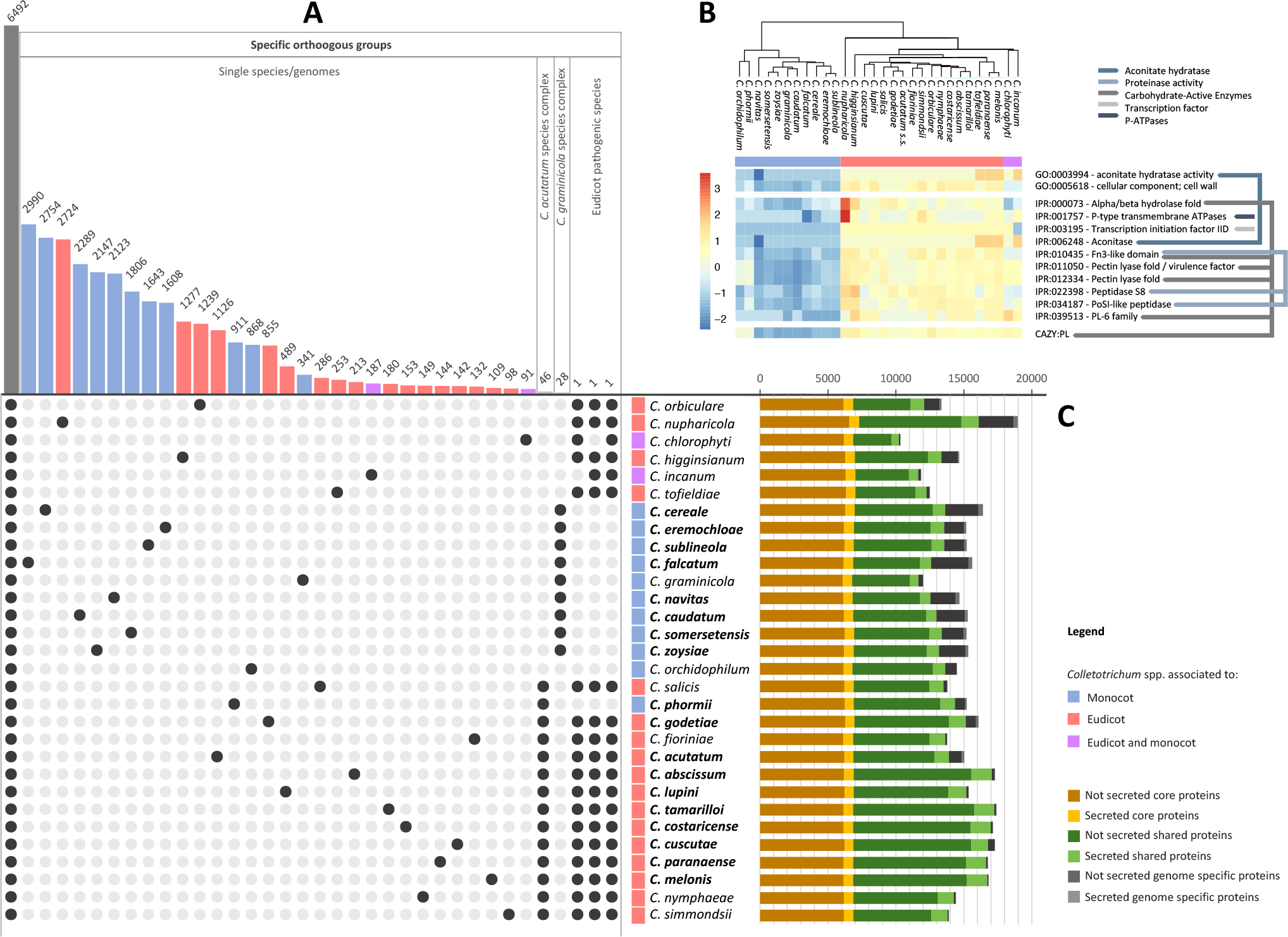
Comparative genomic analysis of *Colletotrichum* species. (**A**) UpsetR plot of the protein clustering analysis. Bars in the upper side represent the number of orthogroups shared by the species highlighted by the black dots reported in the bottom side. (B) Hierarchical clustering of disjoint sets of terms and gene families identified in *Colletotrichum* species associated with monocots and eudicots hosts. Gene Ontology and InterPro terms corresponding to the rows are reported on the right; coloured lines connect overlapping terms. Hierarchical clustering of genes and species was performed and visualized using UPGMA algorithm. Overrepresented (orange to red) and underrepresented functional domains (blue). (**C**) Bar diagrams showing the number of proteins shared with all included species (in yellow), shared with at least two but not all (in green) and those found in only one species (in grey). The light shading indicates for each group the portion of proteins predicted to be secreted.

*Colletotrichum* species diverged from members of the closest related genus *Verticillium* in the late Jurassic around 136.43 million years ago (mya) (186.35 - 99.88) (Supplementary 1). The diversification of species within the genus, based on the estimation of divergence between the two most distantly related species *C. orbiculare* and *C. abscissum*, took place during the Upper (or Late) Cretaceous period, 68.76 mya (103.85 – 45.53). These results suggest that the common ancestor of *Colletotrichum* was associated with eudicots and at least three independent host jumps between eudicots and monocots took place during the evolution of this pathogen. The first took place in the Paleogene (around 25 mya) when species of the Graminicola complex diverged from those belonging to the Spaethianum complex. Interestingly, the diversification of *Colletotrichum* species adapted to plant species belonging to the Poaceae, happened around 20 mya, coinciding with the expansions of grasses from their water-bank habitat into open tracts and their diversification^24^. The second happened in around 15 mya when *C. orchidophilum* diverged from the ancestor of the Acutaum species complex. The third host jump occur in the Neogene around 3.5 mya when the flax pathogenic species *C. phormi* diverged from its closest related species *C. salicis*.

### *Colletotrichum* species associated with monocots have gone through lineage specific expansions of genes and losses of degradative enzymes and other conserved functions

To examine core features shared by all *Colletotrichum* species, by complexes, by individual species, as well as features specific of eudicot and monocot associated species, all predicted proteomes were clustered into groups of orthologous genes (Figure 2A). This approach enabled the identification of the core, shared and species-specific proteins and orthologs only present in species associated with eudicot or monocot hosts.

Enrichment analyses of the core, shared and lineage specific (secreted and non-secreted) genomes did not identify functional category or gene family expansions associated with host range. Considering that the analyses carried out are affected by the sampling, as closely related species are likely to have more shared genes compared to species that are more distant from others, our analyses also highlight that monocot pathogenic species have generally more specific genes compared to eudicot pathogenic species (Figure2A and 2C). While no orthogroups specific to the monocot pathogenic species were identified, we were able to identify three orthogroups only present in those species capable of infecting eudicot plants. These were OG0010350, with one or two copies of the gene present in all eudicot pathogenic species and in *C. incanum* and characterized as a secreted β-glucosidase (CAZy - GH3/FN3), OG0010637 with one or two copies of the gene present in all eudicot pathogenic species and in *C. incanum* and characterized as a secreted protein with unknown function containing a (FAD)-binding domain, and OG0011101 present in all eudicot pathogenic species and in those that have been associated with eudicot and monocot and characterized as an α-1,2-mannosidase (CAZy - GH92). Analyses of functional annotations highlighted two gene ontology (GO), 12 InterPro (IPR) terms and two gene families expanded in eudicot associated species compared to the monocot associated species (Figure2B). No terms were expanded in monocot associated *Colletotrichum* spp. confirming the pattern observed in the analyses based on protein similarity and the two species capable of infecting both hosts (*C. chlorophyti* and *C. incanum*) cluster with the eudicot associated pathogens. As many IPR and GO terms overlap, the results were manually inspected to avoid redundancy.

Overall, terms identified as expanded in eudicot associated pathogens could be clustered into five functional groups (Figure2B): 1) aconitases are genes encoding for enzymes that catalyse the stereo-specific isomerization of citrate to isocitrate in the Krebs cycle; while eudicot pathogens have three or four copies of this gene, monocot pathogens have only two. 2) P-ATPases are proteins that are involved in transport of a variety of different compounds. 3) Transcription initiation factor IID is a general transcription factor (GTF) involved in accurate initiation of transcription by RNA polymerase II. 4) Serine proteases belonging to the MEROPS peptidase family S8. 5) Several terms identified, such as the apha/beta hydrolase fold, the pectin lyase fold, the PL6 family domains as well as others are associated with CAZymes.

To better understand the landscape of CAZymes we focused our analyses on families involved in the degradation of plant polymers. The eudicot infecting species have a higher overall number of genes encoding putative plant biomass degrading enzymes than the species with monocot hosts (Suppl7_CAZYmes_PBD.xlsx), which confirms previous studies^6^. This is also clear by the number of CAZy families encoding carbohydrate esterases (CE), glycoside hydrolases (GH) or polysaccharide lyases (PL), for which the eudicot infecting species have a significantly higher number of genes, even though this difference per family is often small. In contrast, higher gene numbers per family for the monocot infecting species are only present in CE1, GH10, GH11, GH13_1, GH45 and GH62.

Interestingly CE1, GH10, GH11 and GH62 are all involved in xylan degradation, a prominent component of monocot cell walls. In particular, CAZy families encoding putative pectinolytic enzymes have higher numbers of genes in the eudicot infecting species, such as CE8, CE12, GH28, GH43, GH52, GH53, GH78, GH88, GH93, PL1, PL3, PL11 and PL26. However, also CAZy families with putative enzymes targeting lignin (AA1), cellulose (GH1, GH3, GH5, GH7) and hemicellulose (CE16, GH12, GH27, GH36, GH74, GH115) are enriched in the eudicot infecting species. At the individual species level, *C. nupharicola* stands out with an increased number of genes in several CAZy families (AA1_3, CE1, GH1, GH2, GH7, GH28, GH43, GH78). The *Colletotrichum* species lack the subfamily AA1_1 *sensu stricto* laccases but possess putative laccase-like multicopper oxidase encoding genes from the subfamilies AA1_2 and AA1_3. A previously described laccase (*lac2*), which is involved in melanisation in appressorial cells of *C. orbiculare* ^25^, is categorized as a member of family AA1 without a subfamily division, whereas a *C. orbiculare lac1* which does not have a role in melanin biosynthesis or pathogenicity^25^, is catalogued to AA1_3. For three of the species, *C. acutatum*, *C. higginsianum* and *C. graminicola*, growth profiles on plant biomass related substrates are available in the FUNG-GROWTH database^26^. Comparison of the CAZome of these three species (Suppl7_CAZYmes_PBD.xlsx) to their growth profiles did not provide clear correlations. Growth on xylan, galactomannan (guar gum) and inulin is relatively poor for *C. higginsianum* compared to the other two species, but no strong reduction in xylanolytic, mannanolytic or inulinolytic genes can be found in its genome. This evidence also suggests that the CAZyme content in the genome can only partially explain its degradative capability.

To confirm these results and to gain a better understanding on the evolution of the genes identified using both approaches (similarity-based protein clustering and protein terms enrichment) further analyses were carried out. Results of selected CAZy families (GH3 and GH92), aconitases and transcription initiation factors IID (Suppl11_Evo_genefamilies) revealed gene losses in the monocot associated species lineages.

### Transcriptome profiles on different carbon sources reveal a strong variation among species and key genes involved in the interaction with PCW

To identify genes involved in the interaction with the PCW, we performed a transcriptome analysis of four reference species, two eudicot pathogens: *C. higginsianum*, *C. nymphaeae* and two monocot pathogens: *C. graminicola* and *C. phormii*, on three different substrates: glucose, sugar beet pulp (eudicot cell walls: ECW) and maize powder (monocot cell walls: MCW). The four species show different evolutionary histories and genetic distances with *C. phormii* and *C. nymphaeae* being closely related members of the same complex, but associated with monocot and eudicot hosts, respectively.

*C. higginsianum*, *C. phormii* and *C. nymphaeae* have similar patterns of gene expression when the pairwise comparisons of transcriptome profiles are plotted in a principal component analysis (PCA; Figure 3A). In these three species the comparison of genes differentially expressed in ECW vs MCW show a lower diversity compared to the one highlighted in the comparison of genes differentially expressed in both substrates vs. glucose (Figure 3A). This pattern is also confirmed by the overall number of differentially expressed genes (DEGs), where *C. higginsianum*, *C. phormii* and *C. nymphaeae* have the lowest number of both up- and downregulated DEGs in ECW vs MCW while *C. graminicola* has a comparable number of DEGs in other pairwise comparisons (Figure 3B). The differences shown by *C. graminicola* might reflect the longer evolutionary history of association with its host as well as the differences in plant substrate composition between the hosts. Among the four species, *C. nymphaeae* regulate differentially more genes compared to the other species.

**Figure 3.**
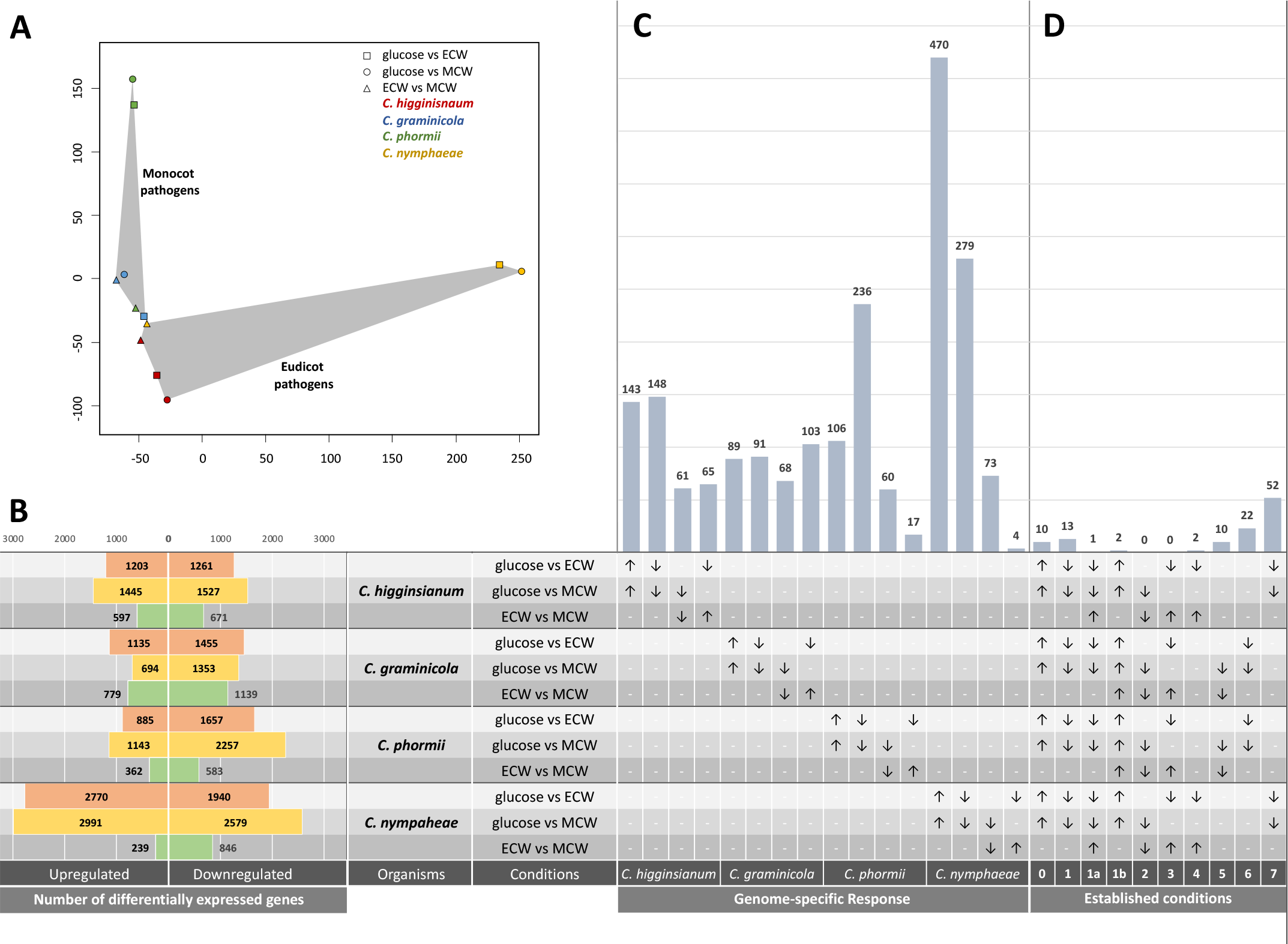
Comparative transcriptomic analysis of selected *Colletotrichum* species (*C. higginsianum*, *C. graminicola*, *C. phormii* and *C. nymphaeae*) on three on different carbon sources: glucose, sugar beet pulp (as eudicot cell walls: ECW) and maize powder (as monocot cell walls: MCW). **(A)** Principal components analysis (PCA) of differentially expressed genes in all conditions analysed; grey areas highlight profiles of monocot and eudicot pathogens. **(B)** Number of differentially expressed genes of each *Colletotrichum* species and in each condition analysed. **(C&D)** Genome specific response represented as the number of genes differentially expressed. For each pairwise comparison and species over expressed genes are indicated by an arrow pointing up while those under expressed are indicated by an arrow pointing down **(C)** Number of genes overexpressed in glucose for each genome are reported in the first column on the left, those overexpressed in PCW are reported in the second column, those overexpressed in MCW are reported in third column and those overexpressed in ECW are reported in forth column. (**D**) Numbers of genes showing the same expression patterns in the intersections described in Table 1.

To better understand the specificity of the response to different substrates, we identified species specific genes overexpressed in presence of glucose, eudicot cell walls (ECW), monocot cell walls (MCW), plant cell walls (PCW: as those genes overexpressed in presence of both ECW and MCW), as well as those shared among all four species, among the eudicot pathogens and among the monocot pathogenic species (Figure 3C and 3D).

Results highlighted a strong specific response by the four species, as the majority of the DEGs are not shared between the four genomes but are specific for each organism.

Comparative analysis of enrichment profiles highlighted five terms enriched among overexpressed genes in eudicot pathogens on ECW (condition 7), all of which (GO:0000981, GO:0006355, IPR001138, IPR036864, PF00172) are associated with Zn(2)-Cys(6) fungal-type DNA-binding domain and transcription regulation. As no other enrichment was identified we performed a manual annotation of genes identified in Figure 3D. As a result, more than one third of all genes identified (32/112) were assigned to three major groups, i.e., transporters, CAZymes and transcription factors.

We identified ten orthologous genes overexpressed in the presence of glucose compared to plant cell walls (condition 0). Among these, four are transporters, three are associated with primary metabolism (such as citrate and fatty acid synthase and sorbitol dehydrogenase), one is a secreted flavoenzyme and two are secreted proteins of unknown function. Sixteen orthologous genes in each species were upregulated in the presence of the substrates containing plant biomass (condition 1, 1A and 1B). In this set we identified four transporters, two transcription factors, three genes belonging to CAZy families GH27, GH5_16 and GH43, and one subclass M28 peptidase. Interestingly one orthogroup (OG_12813) assigned to condition 1a and therefore to genes overexpressed in the presence of plant cell wall by all four species but more overexpressed in eudicot pathogenic species compared to the monocot pathogenic species, has been assigned to the CAZy subfamily GH43 (Table 1) that contains xylan and pectin degrading enzymes.

**Table 1.** Description of transcription profiles, number of genes identified and main biological functions in each condition.

| Condition | Description * | # genes | Main biological function / description |
| --- | --- | --- | --- |
| 0 | genes overexpressed in the presence of glucose | 10 | primary metabolism; transporters |
| 1 | genes overexpressed in the presence of PCW | 13 | CAZy GH27 / GH5 / GH43; transporters; 2 transcription factors |
| 1a | genes overexpressed in the presence of PCW and overexpressed in ECW in eudicot pathogens | 1 | CAZy GH43 |
| 1b | genes overexpressed in the presence of PCW and overexpressed in MCW in monocot pathogens | 2 | Sugar transport; alkaline phosphatases |
| 2 | genes overexpressed in the presence of MCW | 0 | NA |
| 3 | genes overexpressed in the presence of ECW | 0 | NA |
| 4 | genes overexpressed in the presence of ECW only in eudicot pathogens | 2 | CAZy GH142; transmembrane protein |
| 5 | genes overexpressed in the presence of MCW only in monocot pathogens | 10 | CAZy GH11(CBM1); transmembrane proteins; 2 transcription factor |
| 6 | genes overexpressed in the presence of PCW only in monocot pathogens | 22 | CAZy GH43 /GH62(CBM1); transporters, oxidoreductase activity; 3 transcription factors |
| 7 | genes overexpressed in the presence of PCW only in eudicot pathogens | 52 | Unknown functions, Zinc finger – nucleic acid binding; 6 transcription factors |

\* PCW: plant cell wall; MCW: monocot PCW; ECW: eudicot PCW

Two orthogroups were identified as overexpressed in the presence of eudicot cell walls (ECW) only by eudicot pathogens (condition 4) and ten orthogroups were identified as overexpressed in the presence of the monocot cell walls (MCW) only by monocot pathogens (condition 5). This suggests a certain level of specificity by the eudicot and monocot pathogenic species. The main differences between the two sets of genes are the presence of specific transcription factors in the response of the monocot pathogens while the response of the eudicot pathogens lacks specific transcription factors. Another difference is highlighted by differences in genes encoding for CAZy (GH142 in condition 4 and GH11 (CBM1) in condition 5). An opposite situation was observed in condition 6 compared to condition 7, where the number of orthogroups overexpressed by eudicot pathogenic species is more than double of those overexpressed by monocot pathogenic species in the substrates containing plant biomass (MCW or ECW). Both sets are rich in transcription factors, but while *C. graminicola* and *C. phormii* overexpressed several shared genes encoding for CAZymes (such as GH62, AA3_2 and two different genes belonging to the GH43), *C. nymphaeae* and *C. higginsianum* overexpressed only one (also belonging to GH43).

### CAZymes

In contrast to the small differences in gene numbers per CAZy family, comparison of the transcriptome profiles of *C. higginsianum*, *C. nymphaeae*, *C. phormii* and *C. graminicola* revealed high diversity between them. Based on the expression differences of CAZy genes between transcriptome of fungi growth in glucose and other two carbon sources (SBP and MP), the expression of the orthologous genes was clustered for the four fungal species (Figure 4).

**Figure 4.**
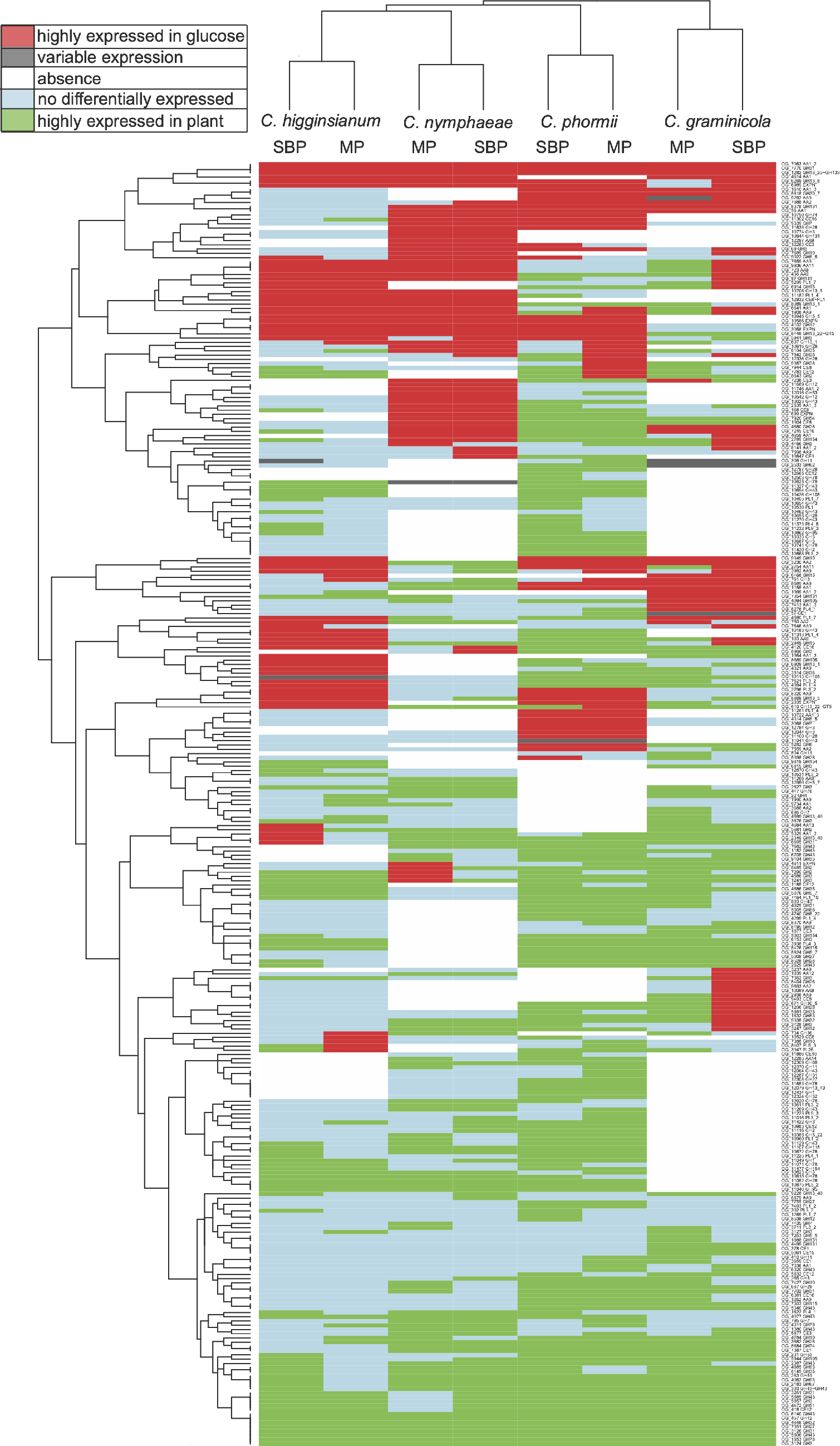
Comparison of differential gene expression of CAZy ortholog groups on PBM or glucose. The genes were binned into the following five categories: highly expressed in glucose, variable expression, absence, no differentially expressed and lowly expressed in glucose for each ortholog gene(s) of each species in each specific comparison and shown in different colours on the figure.

This demonstrated that the transcriptional profiles of the same fungus grown on two different substrates (maize powder and sugar beet pulp) cluster together, indicating that the fungal species had a higher influence on the expression profile than the mono- or eudicot nature of the substrate. The eudicot infecting fungal species (*C. higginsianum*, *C. nymphaeae*) were most similar to each other, while the two monocot infecting species (*C. phormii*, *C. graminicola*) were more distinct. This effect seems to be mainly at the individual orthogroup level, as more similarity can be observed between the fungal species when the number of genes that were upregulated on plant biomass or on glucose were compared between the species for each CAZy family (Figure 5). In this comparison, the clustering of the eudicot fungal infecting species was no longer observed, suggesting strong differences in the transcriptional response of the individual species.

**Figure 5.**
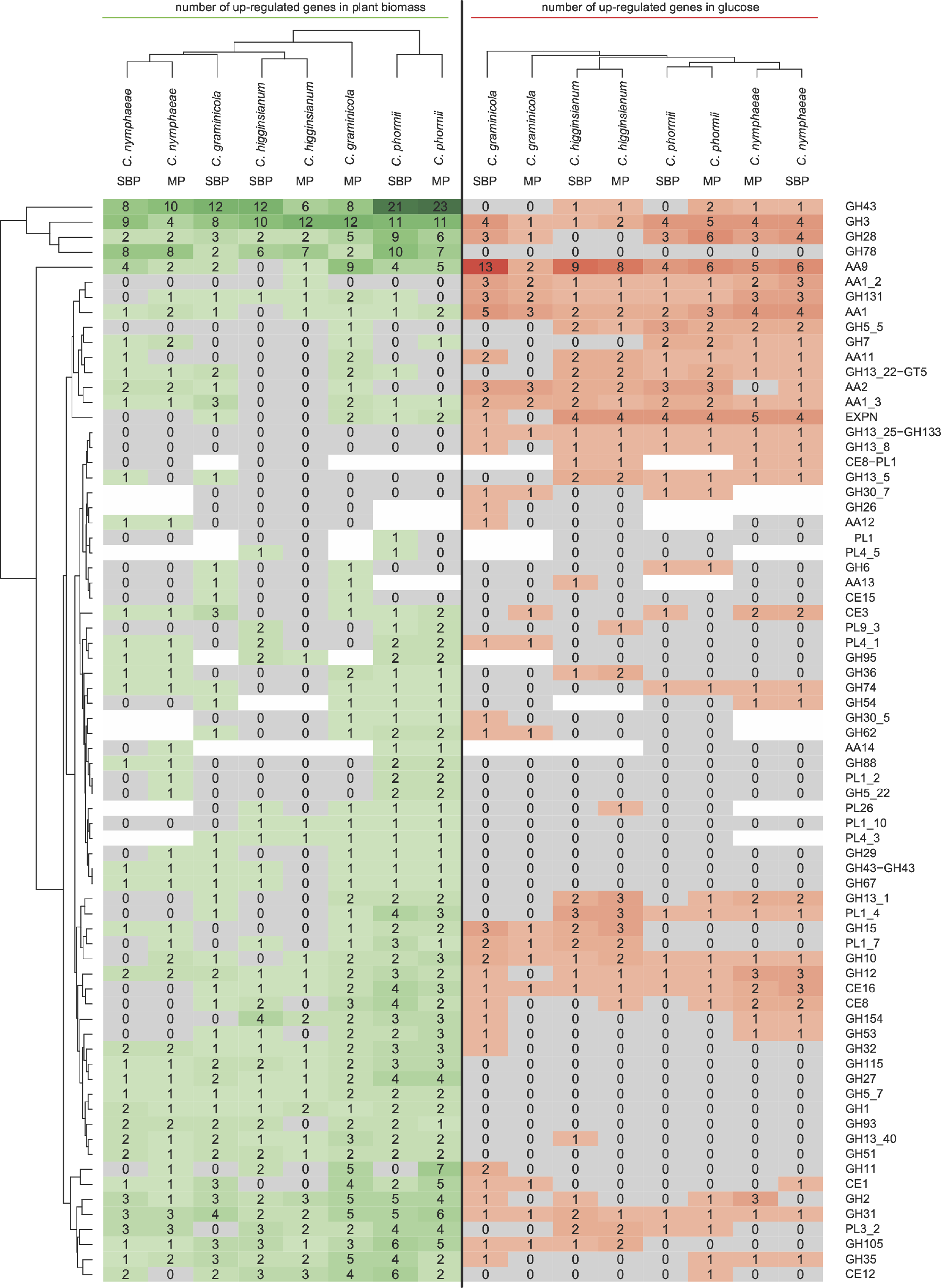
Comparison of the number of CAZy genes upregulated on PBM or glucose for the tested species. The number of highly and lowly expressed genes detected in glucose condition were marked with red and green, respectively. The ortholog genes missed in the specific species were indicated with white colour.

### Transcription factors and gene regulation

The expression profiles of *C. higginsianum*, *C. nymphaeae*, *C. phormii* and *C. graminicola* revealed the presence of several genes encoding transcription factors (TFs) and other regulatory genes showing interesting patterns of expression (Table 2).

**Table 2.** Transcription factors and other genes involved in modulating gene expression identified in the transcriptome dataset.

| Condition | Description * | Orthogroup | Domain | Predicted/putative function |
| --- | --- | --- | --- | --- |
| 1 | genes overexpressed in presence of PCW | OG_1905 | Cys6Zn2 TF | Unknown/vegetative asexual development |
|  |  | OG_7409 | Cys6Zn2 TF | Activator of stress 1 (ASG1)/hyphal growth |
| 5 | genes overexpressed in presence of MCW only in monocot pathogens | OG_8644 | Methyltransferase | Secondary metabolism |
|  |  | OG_1140 | Cys2His2 TF | Unknown |
| 6 | genes overexpressed in presence of PCW only in monocot pathogens | OG_6982 | Methyltransferase | Unknown/putative growth control |
|  |  | OG_401 | Cys6Zn2 TF | Activator of purine utilization |
|  |  | OG_1148 | Cys6Zn2 TF | Secondary metabolism |
| 7 | genes overexpressed in presence of PCW only in eudicot pathogens | OG_2149 | Cys6Zn2 TF | Conidiophore development, hyphal growth |
|  |  | OG_2547 | SFN2 helicase | Chromatin remodeling/DNA repair |
|  |  | OG_3693 | Cys6Zn2 TF | Unknown |
|  |  | OG_2666 | Cys6Zn2 TF | Cutinase transcription factor 1 (CTF1) |
|  |  | OG_2742 | GATA-like TF | Development and disease |
|  |  | OG_5209 | E3 Ubiquitin ligase | Proteasome-mediated ubiquitin-dependent protein catabolic process |
|  |  | OG_7815 | bZIP TF | Oxidative stress/pathogenicity |
|  |  | OG_8935 | GATA TF | Sensing |
|  |  | OG_94 | E3 Ubiquitin ligase | Ubiquitin ligase/histone regulation |

\* PCW: plant cell walls; MCW: monocot cell walls

Surprisingly none of them are orthologs of already characterized TFs directly involved in plant cell walls degradation, some of which are unknown, or we could not identify a clear function. Indeed, all four fungal species overexpressed only two TFs in presence of PCW (condition 1) which have putative function in vegetative and stress growth suggesting that the saprophytic stage of *Colletotrichum* spp. required a re-shaping of the growth *modus operandi*. Interestingly no TFs were overexpressed in the four fungal species growing on MCW (condition 2) or ECW (condition 3), matching with the CAZymes expression profile where species appeared to have a higher influence than the nature of the substrate. Monocot and eudicot associated pathogens responded differently to PCW at the regulatory level. Monocot pathogens specifically overexpressed a narrow set of TFs (five in total), mainly involved in growth control and secondary metabolism. Moreover, only monocot pathogens appeared to be partially adapted to their natural substrate as two TFs are overexpressed in MCW only by monocot pathogenic species (condition 5) while no TFs were differentially expressed in eudicot pathogenic species on ECW. These two TFs show an interesting behaviour: the methyltransferase OG_8644 is present in all four species but differentially expressed only in monocot associated pathogens on MCW, while the unknown Cys2His2 TF OG_1140 is present only in *Colletotrichum* spp. associated with monocot, suggesting that it has been acquired during the adaptation toward monocot host(s)

In contrast to monocot associated pathogens, eudicot pathogens had more expanded and complex regulatory responses with more than half of the total differentially expressed TFs, with no TFs specifically differentially expressed in ECW (condition 4), suggesting that these strains have a less substrate specific response.

Six TFs and three regulatory factors were identified as overexpressed in both substrates (MCW and ECW) only by eudicot associated pathogens (condition 7), although their present in all four genomes. This evidence suggests that these regulatory genes may have lost the function to respond to plant cell walls during the process of host (monocot) adaptation. Most of such TFs appears to have putative function(s) in virulence and pathogenicity. The other regulatory genes found in this category, have general function(s) in chromatin remodelling and/or post transcription regulation, suggesting that the adaptation to eudicot host(s)/substrate(s) also required a fine tune at post-transcriptional/translational level. Confirming this hypothesis, in this category we found several genes involved in translation process/modification, especially at tRNA level (suppl. file 14). This indicating that the chromatin remodeling and the post-translation processes are important for the eudicot associated pathogens for host interactions and/or plant cell wall interaction.

## DISCUSSION

The ancestral *Colletotrichum* was associated with eudicot plants and certain branches progressively adapted to different monocot hosts. The diversification of species inside the genus took place during the Upper (or Late) Cretaceous, 68.76 mya (103.85 – 45.53). This period was characterized by the ecological success of angiosperms that appeared in the fossil records (between 145 and 66 mya)^27^. Previous studies indicate that ancestral angiosperms lived in low evaporative niches during the Early Cretaceous^28^ before the period of their quick diversification in the Mid Cretaceous^29^. During the Late Cretaceous, evolving angiosperms spread towards the poles^30^ and gained ecological dominance in most of the world’s ecosystems by replacing gymnosperms in the evaporatively more demanding upper canopy^31^. In our dataset at least three different events of host jumps and specialization to monocots were detected, the first when species belonging to the Graminicola complex diverged from those belonging to the Spaethianum complex around 25.23 mya (42.71 – 14.91), the second when *C. orchidophilum* diverged from the common ancestor of species belonging to the Acutatum complex around 14.58 mya (23.89 – 8.89) and the third event when *C. phormii* diverged from the closely related species *C. salicis* around 3.45 mya (6.55 – 1.82).

All members of the Graminicola complex are pathogenic to species belonging to the Poaceae. However, while most of the species can infect plants belonging to the Panicoideae subfamily (PACMAD clade), *C. zoysiae* is pathogenic to *Zoysia tenuifolia* which belongs to the Chloridoideae subfamily (PACMAD clade) and *C. cereale* is pathogenic to *Poa annua* which belongs to the Pooideae subfamily (BOP clade). The ancestor of all hosts of the Graminicola species can be placed at the crown node of BOP and PACMAD that is dated at 57 mya (75 – 51 mya) in the late Paleogene^24^. This event happened before the differentiation of species belonging to the Graminicola complex and those belonging to the Spaethianum complex while the quick species diversification into Graminicola species took place between the Miocene and the Oligocene, 18.59 mya (32.56 – 10.62 mya) overlapping with the occupation of open habitats in Africa of their hosts that occurred in the late Eocene–early Oligocene. The Oligocene period was considerably drier than the rest of the Tertiary and these factors might have had an effect on the decrease of the forest cover and the expansion of open habitats^32^. The second jump to monocot hosts happened when *C. orchidophilum* diverged from the ancestor in common with species belonging to the Acutatum complex around 14.58 mya (23.89 – 8.89). *C. orchidophilum* is host specific, infecting different species belonging to the Orchidaceae including species belonging to *Phalaenopsis*, *Cycnoches*, *Dendrobium* and *Vanilla*genera^12, 14, 33^ covering the entire diversity of the Orchidaceae. Previous studies reported that the common ancestor of orchids was supposed to have existed much earlier, between 76 and 84 mya^34^. The last of the three monocot specialization events happened when *C. phormii* diverged from the closely related species *C. salicis* in the Neogene, around 3.45 mya (6.55 – 1.82). *C. phormii* is a worldwide-distributed pathogen of *Phormium* spp. *Dianella*-like fossils from the Eocene have been placed at the crown of the genera *Phormium* and *Dianella*, dating the divergence between these two genera to around 45 mya (SD = 1.0)^35^, which is much earlier than the estimated appearance of *C. phormii*. Among the three events described, *C. orchidophilum* and *C. phormii* might have acquired a key gene or genes that allow the host jump after the appearance of the host while the ancestor of species belonging to the Graminicola complex have evolved simultaneously with its hosts. Interestingly, all lineages of *Colletotrichum* associated with monocots show a certain level of host specificity.

Analyses of functional annotations highlighted gene families expanded in eudicot-associated species compared to the monocot ones. No functional categories were expanded in monocot associated *Colletotrichum* spp. confirming the pattern observed in the analyses based on protein similarity. To better understand the landscape of CAZymes we focused our analyses on the families involved in degradation of plant polymers. Analysis of the plant biomass degradation (PBD) related CAZome of the different *Colletotrichum* species did not reveal large differences, especially when compared to similar studies in the genus *Aspergillus*^36, 37^. The eudicot infecting species have a higher overall number of genes encoding putative plant biomass degrading enzymes than the species with monocot hosts, which confirms results found on a previous study comparing *C. higginsianum* and *C. graminicola* genome^6^. This is also apparent by the number of CAZy families encoding carbohydrate esterases (CE), glycoside hydrolases (GH) or polysaccharide lyases (PL), for which the eudicot infecting species have a significantly higher number of genes, even though this difference per family is often small. In contrast, higher gene numbers per family for the monocot infecting species are involved in xylan degradation, a prominent component of monocot cell walls. This difference between the monocot and eudicot infecting species reflects the more diverse cell walls of eudicots^38^, which would require a broader set of enzymes to efficiently degrade them. A clear difference between monocot and eudicot infecting species was found in the number of genes encoding putative pectin degrading enzymes. Pectin is a major component of eudicot cell walls, but nearly absent in monocots^38^. Studies of specific CAZymes in *Colletotrichum* spp. are relatively few, and they only address some of the enzymes involved in plant biomass degradation. Examples of these are the publication of the crystal structure of a GH28 endopolygalacturonase from *C. lupini*^39^ and the report on polygalacturonase activity for *C. acutatum*^40^. These enzymatic properties cannot be readily attributed to the lifestyle of these plant pathogens. Hydrogen peroxide (H_2_O_2_) may have multiple roles in plant pathogenic fungi because two subfamily AA5_2 alcohol oxidases have been characterized from *C. graminicola* and *C*. *gloeosporioides*^41, 42^. These enzymes have broad substrate ranges and oxidize aliphatic primary alcohols to the corresponding aldehydes, by simultaneously reducing oxygen to hydrogen peroxide. Although their natural substrates have not yet been identified, these enzymes were suggested to have a role in plant cell wall degradation. In addition, an AA5_2 raffinose oxidase that uses trisaccharide raffinose as its preferred substrate, has been characterized from *C. graminicola*^43^.

Moreover, a recent study showed that another AA5_2 paralog from *C. graminicola* oxidizes aryl alcohols to the corresponding aldehydes, thus describing aryl alcohol oxidase activity in the CAZy family AA5, which is traditionally related to AA3 glucose methanol choline (GMC) oxidoreductases^44^. The results of transcriptomic analysis of four reference species on different plant cell wall substrates indicate a strong specific response by each species, as the majority of the differentially expressed genes are not shared between the four species. No genes overexpressed by all four species exclusively in the presence of MCW or ECW were identified. Overall, the results indicate a higher substrate specificity in the monocot pathogenic species *C. graminicola* and *C. phormii* while the response of the eudicot pathogenic ones seem to be more associated with the general presence of plant cell walls. In contrast to the low differences in gene numbers per CAZy family, comparison of the transcriptomes of *C. higginsianum*, *C. nymphaeae*, *C. phormii* and *C. graminicola* revealed high diversity in gene expression. This matches studies on other fungal genera, such as *Aspergillus*, where a proteomic comparison of a large set of species revealed a much higher diversity than was expected based on genome content^45, 46^. These results in part match with previous studies of the production of plant biomass degrading enzymes in *Colletotrichum*. *C. graminicola* has been shown to produce β-glucosidase, β-xylosidase and xylanase activity during solid-state fermentation on different plant biomass substrates . Enzyme families containing these activities (GH1, GH3, GH10, GH11, GH43) were also expressed on plant biomass in our study. Studies into the expression of specific genes revealed monomeric inducers of the responsible regulatory systems. An endopolygalacturonase encoding gene of *C. lindemuthianum* was expressed in the presence of L-arabinose and L-rhamnose^48^. Several of the CAZy genes of *Colletotrichum* have been implicated in pathogenicity , such as an *C. lindemuthianum* endopolygalacturonase encoding gene^49^ and a *C. gloeosporioides* pectate lyase encoding gene (*pel*B)^50^. Transcriptome profiling of *C. graminicola* and *C. higginsianum* has revealed highly dynamic expression of CAZy genes during the infection process. For example, in *C. graminicola*, significant upregulation of several cellulolytic genes, including three GH5, two GH6, three GH7, one GH12, one GH45 and nine AA9 LPMOs, was observed during the necrotrophic phase compared to the biotrophic phase or the formation of the penetration appressorium^3^. In *C. higginsianum*, genes encoding two GH5, one GH6, two GH7 and two GH12 were upregulated during necrotrophic phase compared to *in vitro* growth^6^. In *C. higginsianum* and *C. graminicola*, an orthologous GH131 encoding gene was highly upregulated during both biotrophic and necrotrophic phases, whereas in *C. higginsianum*, another GH131 family gene was also upregulated during appressorial penetration and biotrophic phase^51^. In addition, the corresponding recombinant GH131 proteins were demonstrated to have broad specificity towards substrates with β-1,3- and β-1,4-glucosidic linkages, and they were suggested to have a role in plant cell walls remodeling, breakdown of hemicellulosic structures or facilitating other enzymes to access cellulose^51^. In *C. fructicola*, a transcriptomic study of four types of infection-related structures (conidia, appressoria, cellophane infectious hyphae and infected plant tissue) revealed an upregulated expression of 27 CAZy genes during appressorium formation^52^. Among these genes, 14 encode for redox enzymes with the highest enrichments from AA2 (heme-containing peroxidases) and AA5 (copper radical oxidases, CROs). Under cellophane infectious hyphae, high expression of GH7, AA9, PL1 and CBM1 family members was also detected. As in our study only a single time point was analyzed, this could explain the absence of the induction of some of these genes in our results.

Previous studies have reported gene duplications within this gene family in species characterized by a broad host range^21, 22^. Interestingly, different members of the CAZy family GH43 have been identified in three different conditions. Both of these results suggest that the GH43 may be an important family for plant substrate interaction and/or degradation in *Colletotrichum* species.

The expression of the transcription factors (TFs) and other regulatory genes of *C. higginsianum*, *C. nymphaeae*, *C. phormii* and *C. graminicola* were analyzed based on the orthogroups clustering and according to the different conditions. Unexpectedly, none of the major known TFs involved in plant biomass utilization^2^ passed our requirements/cut off, while most of differentially expressed regulatory genes identified were TFs with uncharacterized function or other regulatory factors, mainly involved in chromatin remodeling. We found differentially expressed TFs specific to plant cell walls, specific to monocot pathogenic species and specific to eudicot pathogenic species on both monocot and eudicot substrates. We did not identify differentially expressed TFs selective for eudicot cell walls, suggesting that the type of substrate (which significantly differ in composition) is an irrelevant factor for eudicot pathogenic species. Exceptions are monocot associated pathogens which overexpressed one TF and one methyltransferase in the monocot cell walls, and two TFs and one methyltransferase in both substrates. This suggests that adaptation to monocots required changes not only at transcriptional level, but also at the chromatin access level. However, half of such TFs and other regulatory factors were overexpressed only by eudicot pathogens, suggesting that eudicot pathogens have a more complex regulation, most likely reflecting the substrate complexity of their host plants.

Overall, these results reveal a moderate level of genomic diversity within the genus *Colletotrichum* with respect to plant biomass degradation, mainly separating monocot and eudicot pathogens. However, a much stronger level of diversity appears to occur at the post-genomic level. This can in part be assigned to the use of non-orthologous members of the same CAZy family by different *Colletotrichum* species. Our results indicate a higher substrate specificity in the monocot pathogenic species *C. graminicola* and *C. phormii* while the response of the eudicot pathogenic species seem to be more associated with the general presence of plant cell walls.

## MATERIALS AND METHODS

### Strains used in this study

The genomes of 18 *Colletotrichum* species were sequenced and compared to the genomes of publicly available representative species (Table3).

### Nucleic acid purification

#### DNA purification for PacBio sequencing

Total genomic DNA was extracted using a SDS-CTAB method^53^ with some modifications. 200 mg of mycelium were placed into a 2 mL sterile extraction tube prefilled with 0.35 g of acid washed silica glass beads (0.5 mm) (Benchmark Scientific Inc., NJ). 50 mg of PVP40 (Sigma-Aldrich, Saint Louis, USA) and 400 µL of ice-cold lysis buffer (150 mM NaCl, 50 mM EDTA, 10 mM Tris-HCl pH 7.4, 30 µg mL^-1^ Proteinase K) were added to the extraction tubes prior homogenization.

The mycelium was homogenized using the bead-beating method through a BeadBug™ Microtube Homogenizer (Benchmark Scientific Inc., NJ). Three cycles of 30 s and 4000 rpm each, were followed by 30 s interval during which the samples were placed on ice.

Sodium dodecyl sulfate (SDS; Sigma-Aldrich, Saint Louis, USA) was added to a final concentration of 2% (w/v) and the mixture was incubated in a water bath at 65°C for 40 min. The lysed mixture was subsequently centrifuged for 10 min at 4°C and 2500 × *g*. The supernatant was transferred to a new tube and the volume was measured to adjust the NaCl concentration to 1.4 M and 1/10 volume of a 10% cetyltrimethylammonium-bromide (CTAB) buffer (10% CTAB, 500 mM Tris-HCl, 100 mM EDTA, pH 8.0) was added. After thorough mixing, the solution was incubated at 65°C for 10 min and cooled at 15°C for 2 min.

An equal volume of a solution of chloroform-isoamyl alcohol (24:1 v/v) was added to the mixture that was then centrifuged for 10 min at 4°C and 6700 × *g*.

The supernatant was transferred to a new tube and the DNA was precipitated with two volumes of 95% cold ethanol. Samples were stored at –20°C for a minimum of 1 hour and subsequently centrifuged 3 min at 4°C and 12,000 × *g*. The resulting pellet was rinsed once with 70% cold ethanol, vacuum-dried and dissolved in nuclease-free water (Promega, Madison, WI, USA). DNA solutions were stored at –20°C until use.

#### DNA purification for Illumina sequencing

Genomic DNA was extracted based on a modified CTAB procedure^54^. The mycelium (250 mg) was ground under liquid nitrogen using a sterilized chilled mortar and pestle. The resultant powder was mixed with 15 ml of a preheated solution (60°C) containing 10% CTAB, 2 M Tris-CI (pH 8.0), 0,5 M EDTA, 1.4 M NaCI and 0.5% 2-mercaptoethanol. After incubation for 30 min at 60°C, samples were washed twice with a 15 ml volume of chloroform:isoamyl alcohol 24:1 (v/v). The aqueous phase was transferred to a clean tube, and the nucleic acids were precipitated with 0.6 volume of cold 2-propanol. After 2-hour incubation at room temperature, the samples were centrifuged for 2 min at 460 × *g*. The pellet was washed twice with 66% (v/v) EtOH and 34 % of 0.1 M NaCl. Tubes were centrifuged at 1500 × *g* for 10 min, washing buffer (supernatant) was removed and pellets were air dried in a fume hood (approximately 1 h). The pellets were resuspended in one ml of AB, left for few minutes, centrifuged for 5 min and supernatant (DNA) saved and pellet discarded.

#### RNA purification

A transfer experiment was performed for transcriptomics. 250 mL of complete medium (CM)^55^ containing 2% D-fructose in 1 L Erlenmeyer flasks was inoculated with 2.5 × 10^8^ fresh spores, harvested from a MEA plate, and incubated in a rotatory shaker at 25°C for 20 h at 140 rpm. The mycelium was harvested by filtration, washed with liquid MM^55^ (without carbon source) and 2.5 g mycelium (wet weight) was transferred to 125 mL Erlenmeyer flasks containing 25 mL MM with 1% of maize powder (MS) or sugar beet pulp (ES), and incubated in a rotatory shaker at 25°C and 140 rpm. After pre-culturing and after 96 h of incubation in MCW or ECW, the mycelium was harvested by vacuum filtration, dried between tissue paper, directly frozen in liquid nitrogen and stored at −80°C^56^. All experiments were performed in triplicate.

### Genome sequencing and assembly

Five strains were sequenced using PacBio reads, namely, *C. godetiae* CBS 193.32, *C. acutatum* CBS 112980, *C. phormii* CBS 102054, *C. lupini* CBS 109225, and *C. navitas* CBS 125086. Genomic DNA was sheared into fragments larger than 10 kb using a Covaris g-TUBE. The sheared DNA was treated with DNA damage repair mix followed by end repair and ligation of blunt adapters using SMRTbell Template Prep Kit 1.0 (Pacific Biosciences). The library was purified with AMPure PB beads. PacBio Sequencing primer was then annealed to the SMRTbell template library and Version P6 sequencing polymerase was bound to them. The prepared SMRTbell template libraries were then sequenced on a Pacific Biosciences RSII sequencer using Version C4 chemistry and 1×240 sequencing movie run times. The filtered subread data was assembled using Falcon version 0.2.2 (https://github.com/PacificBiosciences/FALCON), improved with finisherSC version 2.0, and polished with Quiver version smrtanalysis_2.3.0.140936.p5 (https://github.com/PacificBiosciences/GenomicConsensus).

For the other seven strains *(C. cereale, C. eremochloae, C. sublineola, C. falcatum, C. caudatum, C. somersetensis, C. zoysiae),* genomic DNA was sheared to 300 bp using the Covaris LE220-Plus and size selected with SPRI using TotalPure NGS beads (Omega Bio-tek). The fragments were treated with end-repair, A-tailing, and ligation of Illumina compatible adapters (IDT, Inc) using the KAPA-HyperPrep kit (KAPA Biosystems). The prepared libraries were quantified using KAPA Biosystems’ next-generation sequencing library qPCR kit and run on a Roche LightCycler 480 real-time PCR instrument. The quantified libraries were then prepared for sequencing on the Illumina HiSeq sequencing platform utilizing a TruSeq paired-end cluster kit, v4. Sequencing of the flowcell was performed on the Illumina HiSeq2500 sequencer using HiSeq TruSeq SBS sequencing kits, v4, following a 2×150 indexed run recipe. Raw reads filtered for artifact and process contamination were assembled with Velvet^58^. The resulting assembly was used to create a long mate-pair library with insert 3000 +/- 300 bp which was then assembled with the original Illumina library with AllPathsLG release version R49403^59^.

For transcriptomes, stranded cDNA libraries were generated using the Illumina Truseq Stranded mRNA Library Prep kit. mRNA was purified from 1 μ g of total RNA using magnetic beads containing poly-T oligos. mRNA was fragmented and reversed transcribed using random hexamers and SSII (Invitrogen) followed by second strand synthesis. The fragmented cDNA was treated with end-pair, A-tailing, adapter ligation, and 8-10 cycles of PCR. The prepared libraries were quantified using KAPA Biosystems’ next-generation sequencing library qPCR kit and run on a Roche LightCycler 480 real-time PCR instrument. The quantified libraries were then prepared for sequencing on the Illumina HiSeq sequencing platform utilizing a TruSeq paired-end cluster kit, v4. Sequencing of the flowcell was performed on the Illumina HiSeq2500 sequencer using HiSeq TruSeq SBS sequencing kits, v4, following a 2×150 indexed run recipe. RNA-Seq raw reads were assembled into consensus sequences using either Rnnotator v3.3.2^60^ *(C. eremochloae, C. sublineola, C. falcatum, C. somersetensis, C. zoysiae*) or Trinity ver. 2.1.1^61^ *(C.cereale, C. navitas, C. caudatum, C. godetiae, C. phormii, C. acutatum s.s.* and *C.lupini*). *C. abscissum*, *C. cuscutae*, *C. tamarilloi*, *C paranaense*, *C. costaricense*, *C. melonis* were sequenced using Illumina HiSeq 2500 Rapid-PE 250bp sequencing technology by the McGill University and Genome, Quebec Innovation Centre (Canada). Paired reads were assembled using SPAdes v3.8.2^62^. Scaffolds were filtered for errors and contaminations based on low coverage value. High coverage fragments were manually checked by blastn against the nr database and scaffolds belonging to the mitochondrial genome and to the ribosomal cluster were masked.

### Analysis of genome completeness

BUSCO v3^63^ (Benchmarking Universal Single-Copy Orthologs) was used to search the selected genomes for 3725 Sordariomycete orthologous genes (*sordariomyceta_odb9* data set) to assess the completeness of the genome sequences.

### Gene annotation

The genome sequences of *C. cereale*, *C. eremochloae*, *C. sublineola*, *C. falcatum*, *C. navitas*, *C. caudatum*, *C. somersetensis*, *C. zoysiae*, *C. godetiae*, *C. phormii*, *C. acutatum s.s.* and *C. lupini* were annotated using the JGI annotation pipeline^64^.

The MAKER2 v2.31.8 annotation pipeline^65^ was used to annotate the genome of *C. abscissum*, *C. cuscutae*, *C. tamarilloi*, *C paranaense*, *C. costaricense*, *C. melonis* as previously described^21^, all proteins and transcripts from the *Colletotrichum* ssp. analyzed in this study were used as supporting evidence for gene model annotation.

### Phylogeny and divergence date estimation

The proteomes were clustered with OrthoFinder v0.4^66^ and the clusters were analyzed with Mirlo (https://github.com/mthon/mirlo) to identify the 500 most phylogenetically informative single copy gene families. The families were aligned with MAFFT 7^67^ and then concatenated. A substitution model and its parameter values were selected using ProtTest 3.4^68^. A phylogenetic tree was reconstructed using Bayesian MCMC analysis from the concatenated alignment of the 500 genes identified with Mirlo under the WAG + I evolutionary model and the gamma distribution calculated using four rate categories and homogeneous rates across the tree. The posterior probability threshold was 50%.

A selection of 126 genomes covering the Pezizomycotina plus the genome of *Saccharomyces cerevisiae* as an outgroup were selected from the MycoCosm database (Supplementary 1) and analyzed. The calibrated tree was inferred by applying the RelTime method^69, 70^ to the supplied phylogenetic tree whose branch lengths were calculated using the Ordinary Least Squares method using MEGA X v10.1.7^71^.

The timetree was computed using 5 calibration point (3 fossil records and 2 estimated constraints):

1: Paleopyrenomycites on the crown group of Pezizomycotina, thus assuming the common ancestor of all filamentous, sporocarp-producing Ascomycota (Pezizomycotina) to be 400 mya^72–74^ (normal distribution; standard deviation = 150).
2: *Aspergillus collembolorum*re presenting the common ancestor of the genus *Aspergillus* was constrained to an age of 35 mya^75^ (normal distribution; standard deviation = 15)
3: The fossil Metacapnodiaceae^76^ representing the common ancestor of the order Capnodiales was constrained to an age of 100 mya (normal distribution; standard deviation = 50)
4: Sordariomycetes crown to an age of 207-339 mya^77^ (equal distribution)
5: *Cordyceps* - *Metarhizium* divergence to an age of 146-206 mya^78^ (equal distribution)

The Tao method was used to set minimum and maximum time boundaries on nodes for which calibration densities were provided^79^. The evolutionary distances were computed using the Poisson correction method^80^ and are in the units of the number of amino acid substitutions per site. This analysis involved 127 amino acid sequences and total of 124023 positions in the final dataset.

Evolutionary analyses were conducted in MEGA X^71^.

### Annotation and evolution of specific gene categories

Proteins that are transported out of the cell and into the extracellular space were identified with SignalP-4.1^81^. Protein domains were annotated using Pfam^82^ and InterPro^83^ and mapped to Gene Ontology terms GO terms^84^, CAZymes were annotated using CAZy pipeline^85^.

Peptidases were annotated with the MEROPS database, a hierarchical, structure based classification for peptidases, organized into families and clans (https://www.ebi.ac.uk/merops/)^86^.

BLASTp^87^ and RunIprScan (http://michaelrthon.com/runiprscan/) results were used to manually identify genes encoding enzymes that are signatures of backbone secondary metabolite (SM) genes in the Ascomycota^88^: nonribosomal peptide synthetases (NRPS; IPR010071, IPR006163, IPR001242), polyketide synthases (PKS; IPR013968), DMATS-family aromatic prenyltransferases (IPR017795, Pfam PF11991), and terpene synthases/cyclases (IPR008949).

Transcription factors were identified using BLASTp against NCBI non-redundant protein sequences (nr) database and the Aspergillus Genome Database (AspGD)^89^. P value of 1e-10 was used as cutoff in both cases. NCBI conserved Domains Database(CCD) and EMBL Simple Modular Architecture Research Tool (SMART) (https://smart.embl.de)^90^ were used to manually assign putative function(s) to uncharacterized transcription factors.

Cys_6_Zn_2_ and Cys_2_His_2_ regulators were also analyzed by phylogenetic analyses (NJ) using orthologs of all kingdoms of known regulators involved in plant biomass degradation^2^.

### Comparative genomics

#### Ortholog identification and protein cluster analyses

The Markov Cluster algorithm (mcl v14-137^91^) was used for the identification of protein clusters while (Co-)orthologous groups were identified by Proteinortho v5.16b^92^.

#### The pan- and core-genome and lineage specific genes

1 - The pan-genome: all genes present in one or more species.
2 - The core-genome: genes present in all included species, including paralogs. This set is expected to encode cellular functions needed for all species.
3 – Lineage specific genes: genes found in only one species or cluster in our analysis, with or without paralogs (Included in these, we would expect to find genes involved in environmental adaptation. This group can also include annotation errors)

#### Identification of expansions and contractions of gene families associated with PCW

Functional categories associated with mono- or eudicot pathogenic species were identified using two different statistical analyses.

Disjoint sets calculated as:

Set 1 = monocot pathogens
Set 2 = eudicot pathogens
if (Min Set1 > Max Set2) than term is overrepresented in Set1
if (Min Set2 > Max Set1) than term is overrepresented in Set2

Terms enriched based on Fisher’s exact test were calculated for each in each genome in the following subset: secretomes, all core proteins, secreted core proteins, all shared proteins, secreted shared proteins, all species-specific proteins, and secreted species-specific proteins. Profiles were compared to identify terms enriched only in monocot or eudicot pathogens.

### Transcription profiles of *Colletotrichum* spp. on monocot and eudicot plant cell walls

The CAZy families related to plant biomass degradation were selected based on a previous study^93^. We compared gene expression of the selected CAZy genes between transcriptome of fungi growth on glucose and the other two carbon sources (substrates containing different plant cell walls). The expression difference was binned into the following five categories: highly expressed in glucose, variable expression, absence, no differentially expressed and lowly expressed in glucose for each ortholog gene(s) of each species in each specific comparison. The variable expression means more than one gene were included in the ortholog group of a specific species and the expression of these genes show different expression trend in comparison. Only all the orthologs of a species show the same highly or lowly expression in glucose were defined as differentially expressed gene. The no differentially expression means the gene expression show no significant difference between glucose and other two carbon sources. The absence means the ortholog gene were not detected for specific species. These categories were coded with −2, −1, 0, 1, 2 respectively, and were visualized with heatmap using R package “gplots”, with the complete-linkage clustering method and Euclidean distance. For each CAZy family, we did similar analysis. Instead of summarizing the expression difference between glucose and other carbon sources to different categories, the numbers of genes of highly or lowly expressed in glucose condition were visualized in the heatmap. To distinguish the numbers of highly and lowly expressed gene observed in comparison of transcriptome of fungi growth in glucose and other two carbon sources, we transferred numbers of lowly expressed genes to the corresponding negative values in clustering calculation.

### Identification and analysis of differential gene expression

Stranded RNASeq library(s) were created and quantified by qPCR as described earlier. Sequencing was performed using Illumina HiSeq2500 following a 2×100 indexed run recipe. Raw fastq file reads were filtered and trimmed for quality and contamination. Filtered RNA-Seq reads from each library were aligned to the corresponding reference genome using HISAT version 0.1.4-beta^94^. featureCounts^95^ was used to generate the raw gene counts using genome annotations. Only primary hits assigned to the reverse strand were included in the raw gene counts (-s 2 -p --primary options). DESeq2 version 1.10.0^96^ was subsequently used to determine which genes were differentially expressed between pairs of conditions. The parameters used to call a gene differentially expressed between conditions were log2FoldChange > 2 and p-value < 0.05.

### Comparative transcriptomics

A custom script *orthoexpress.pl* was developed to combine Proteinortho v5.16b^92^ and expression profiles in order to identify groups of genes showing specific expression profiles.

Recent duplications were manually checked. In case of different behavior of paralogs both forms of the (co-)orthologous groups were analyzed independently.

Seven logical conditions (Table 4) were established to identify genes differentially expressed in specific organisms/conditions.

**Table 3.**
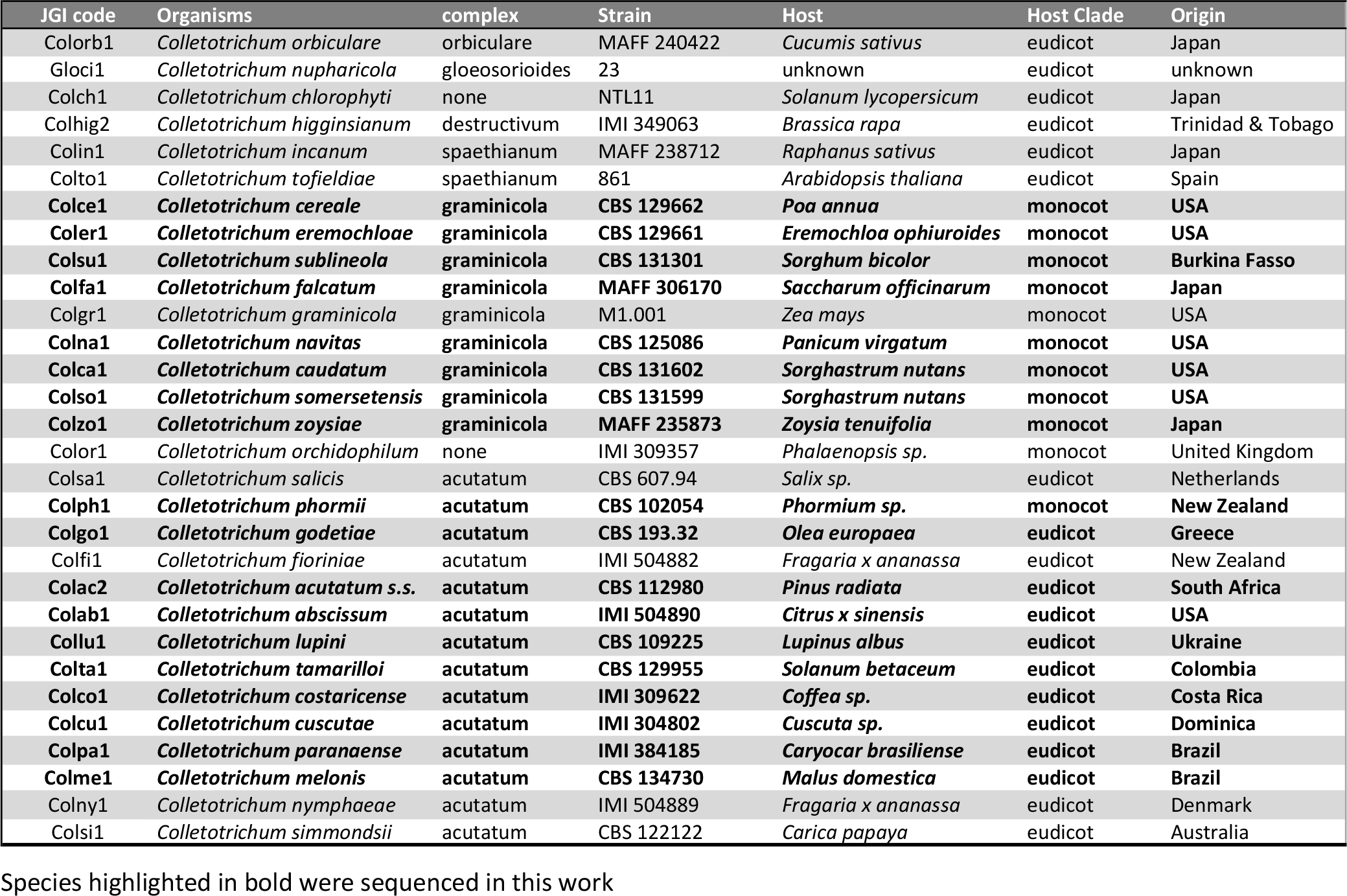
*Colletotrichum* spp. genomes used in this study.

**Table 4.**
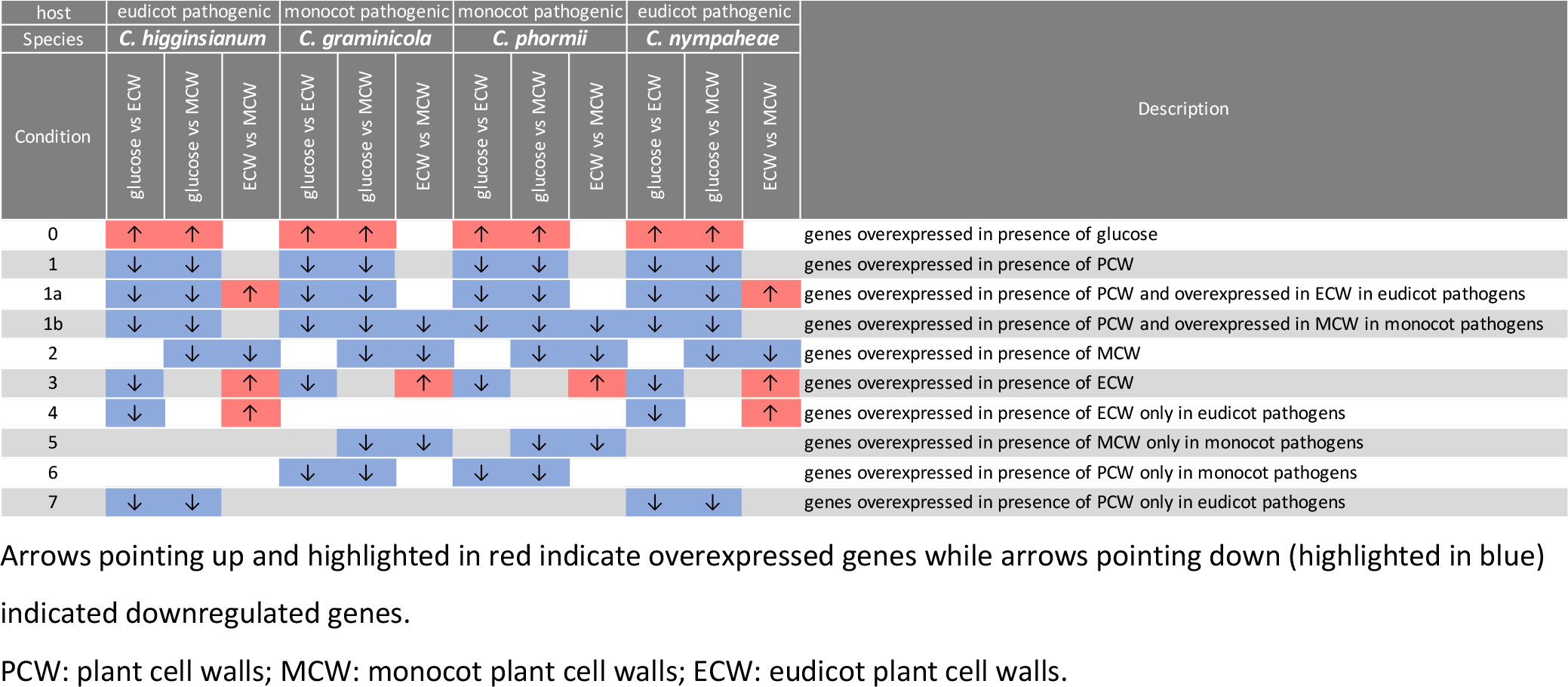
Conditions established for the identification of specific differentially expressed genes

Description of the established conditions of transcription profiles are reported below:

◦ Condition 0: genes overexpressed in presence of glucose
◦ Condition 1: genes overexpressed in presence of plant cell walls (PCW)
◦ Condition 1a: genes overexpressed in presence of PCW and overexpressed in eudicot cell walls (ECW) in eudicot pathogens
◦ Condition 1b: genes overexpressed in presence of PCW and overexpressed in monocot cell walls (MCW) in monocot pathogens
◦ Condition 2: genes overexpressed in presence of MCW
◦ Condition 3: genes overexpressed in presence of ECW
◦ Condition 4: genes overexpressed in presence of ECW only in eudicot pathogens
◦ Condition 5: genes overexpressed in presence of MCW only in monocot pathogens
◦ Condition 6: genes overexpressed in presence of PCW only in monocot pathogens
◦ Condition 7: genes overexpressed in presence of PCW only in eudicot pathogens

## DATA ACCESS

Genome assembly and annotations are available at the JGI fungal genome portal MycoCosm^64^ and are available at DDBJ/EMBL/GenBank. Summary statistics for each genome sequence are reported in Suppl2_Data_access.xlsx.

## ADDITIONAL FILES

**Supplementary File S1:** Time-calibrated phylogenomic tree of 123 fungal genomes belonging to the Pezizomycotina subdivision; *Saccharomyces cerevisiae* genome was used as outgroup. Bars around each node represent 95% confidence intervals. The timetree was computed using 5 calibration points highlighted with red dots (1, 2 and 3 are fossils and 4 and 5 are estimated constraints); see details in the materials and methods section. Major taxonomic classes and respective crown divergent times are reported in green while the crown of *Colletotrichum* is highlighted in orange. Genome information and references are reported in the table below the tree and the genomes sequenced in this work are in bold. Mya = million years ago

**Supplementary Table S2:** Genomes used in this study and relative information.

**Supplementary Table S3:** Gene Ontology (GO) enrichment analysis. For each genome the number of encoded proteins associated with a specific GO term is reported. Statistical comparison of the gene number differences in each GO terms between monocot and eudicot infecting species were compared with the Wilcoxon rank-sum test.

**Supplementary Table S4:** InterPro (IPR) enrichment analysis. For each genome the number of encoded proteins associated with a specific IPR term is reported. Statistical comparison of the gene number differences in each IPR terms between monocot and eudicot infecting species were compared with the Wilcoxon rank-sum test.

**Supplementary Table S5:** Pfam protein families enrichment analysis. For each genome the number of encoded proteins associated with a specific Pfam term is reported. Statistical comparison of the gene number differences in each Pfam terms between monocot and eudicot plant infecting species were compared with the Wilcoxon rank-sum test.

**Supplementary Table S6:** Carbohydrate-Active enZYmes (CAZy) encoding gene enrichment analysis. For each genome the number of encoded CAZy is reported. Statistical comparison of the gene number differences in each CAZy families between monocot and eudicot infecting species were compared with the Wilcoxon rank-sum test.

**Supplementary Table S7:** Comparison of the genome content of 30 *Colletotrichum* species with respect to putative genes involved in plant biomass degradation.

Overall comparison of the species with respect to relevant CAZy families. Statistical comparison of the gene number differences in each CAZy family between monocot and eudicot infecting species were compared with the Wilcoxon rank-sum test.

Differences with a p-value <0.01 are highlighted in yellow. Conditional formatting was applied to the gene numbers in the CAZy family to visualize the differences.

MCO = multicopper oxidase, CDH = cellobiose dehydrogenase, GMC = glucose-methanol-choline oxidoreductase, LPMO = lytic polysaccharide monooxygenases, AXE = acetyl xylan esterase, FAE = feruloyl esterase, PME = pectin methyl esterase,

RGAE = rhamnogalacturonan acetyl esterase, GE = glucuronoyl esterase, HAE = hemicellulose acetyl esterase, BGL = β-glucosidase, MND = β-mannosidase, LAC = β-galactosidase, GUS = β-glucuronidase, BXL = β-xylosidase, EGL = endoglucanase, MAN = endomannanase,

CBH = cellobiohydrolase, XLN = endoxylanase, XEG = xyloglucanase, AMY = α-amylase, AGD = α- glucosidase, GLA = glucoamylase, AGL = α-galactosidase, PGA = endopolygalacturonase, PGX = exopolygalacturonase, RHG = endorhamnogalacturonase, RGX = exorhamnogalacturonase,

XGH = xylogalacturonase, AFC = α-fucosidase, XBH = xylobiohydrolase, AXL = α-xylosidase, INV = invertase, INU = endoinulinase, INX = exoinulinase, ABF = α-arabinofuranosidase, ABN = endoarabinanase, GAL = endogalactanase, AXH = arabinoxylan arabinofuranohydrolase,

AGU = α-glucuronidase, RHA = α-rhamnosidase, UGH = unsaturated galacturonan hydrolase, ABX = exoarabinanase, URGH = unsaturated rhamnogalacturonan hydrolase, AMG = amylo-α-1,6- glucosidase, PLY = pectate lyase, PEL = pectin lyase, RGL = rhamnogalacturonan lyase

**Supplementary Table S8:** Comparison of the gene content of 30 *Colletotrichum* species with respect to putative peptidases and their inhibitors.

**Supplementary Table S9:** Comparison of the gene content of 30 *Colletotrichum* species with respect to putative transporters.

**Supplementary Table S10:** Comparison of the gene content of 30 *Colletotrichum* species with respect to putative transcription factors. Statistical comparison of the gene number differences in each transcription factors families terms between monocot and eudicot plant infecting species were compared with the Wilcoxon rank-sum test.

**Supplementary Figures S11:** Phylogenetic tree of selected gene families based on InterPro (IPR) domain distribution: PL-6 family - IPR039513; Transcription initiation factor IID, subunit 13 - IPR003195; Aconitase, mitochondrial-like - IPR006248; PoSI-like peptidase domain - IPR034187. Red taxa indicate eudicot pathogenic species, blue indicate monocot pathogenic species and purple taxa indicate *Colletotrichum* species that can infect both plant hosts. Pink boxes indicate gene lineages specific of the eudicot pathogens. Number next to the nodes represent support values expressed as % while thicker branches indicate a support value of 100%.

**Supplementary Figure S12:** Correlation matrix of 9 RNA-seq libraries. Pairwise Pearson correlation coefficients (PCC) were calculated for comparison among transcriptomes of various combinations of *Colletotrichum* spp. and substrates. Samples were hierarchically clustered with the Euclidean distance method. The color scale indicates the degree of correlation.

**Supplementary Figure S13:** Volcano plots showing for each pairwise comparison analyzed the genes considered differentially expressed (green dots) based on log2FoldChange > 2 and p-value < 0.05 **Supplementary Table S14:** List of orthogroups and main biological functions related to the genes identified based on specific expression patterns in each condition established. Information such as: foldchange (positive values indicating overexpressed genes are highlighted in red while negative values indicating down regulated genes are highlighted in blue), conserved domains, gene families and locus tags are also reported.

## ACKNOWLEDGMENTS

This research was supported by funds from the Ministerio de Ciencia Innovación y Universidades of Spain (AGL2015-66362-R) and grants RTI2018-093611-B-I00 and PID2021-125349NB-100 from the Ministerio de Ciencia e Innovación (MCIN) of Spain AEI/10.13039/501100011033 and the European Regional Development Fund (ERDF). The work (proposals: 10.46936/10.25585/60000617 and 10.46936/10.25585/60000725) conducted by the U.S. Department of Energy Joint Genome Institute (https://ror.org/04xm1d337), a DOE Office of Science User Facility, is supported by the Office of Science of the U.S. Department of Energy operated under Contract No. DE-AC02-05CH11231. EB was supported by a grant of the Dutch Technology Foundation STW, Applied Science division of NWO, and the Technology Program of the Ministry of Economic Affairs 016.130.609 to RPdV. The Academy of Finland grant number 308284 to MRM is acknowledged. R.B. was supported in part by the postdoctoral program of USAL (Programme II).

We would like to thank the staff at the Plataforma Andaluza de Bioinformática of the University of Málaga, Spain, for providing computer resources and technical support.

The authors would also like to thank Francis Martin and Rytas Vilgalys for the permission to use the genome of *Glomerella cingulata* 23 (= *Colletotrichum nupharicola* 23); Jon Magnuson for the permission to use the genome of *Sclerophora sanguinea* CBS 100924; Olafur Andresson for the permission to use the genome of *Lobaria pulmonaria* Scotland reference genome; Dave Greenshields for the permission to use the genome of *Penicillium fellutanum* ATCC 48694.

## Notes

### Competing Interest Statement

The authors have declared no competing interest.

