## Supplementary material for "Genome evolution and transcriptome plasticity associated with adaptation to monocot and eudicot plants in *Colletotrichum* fungi": Suppl1_Calibrated_tree.pdf

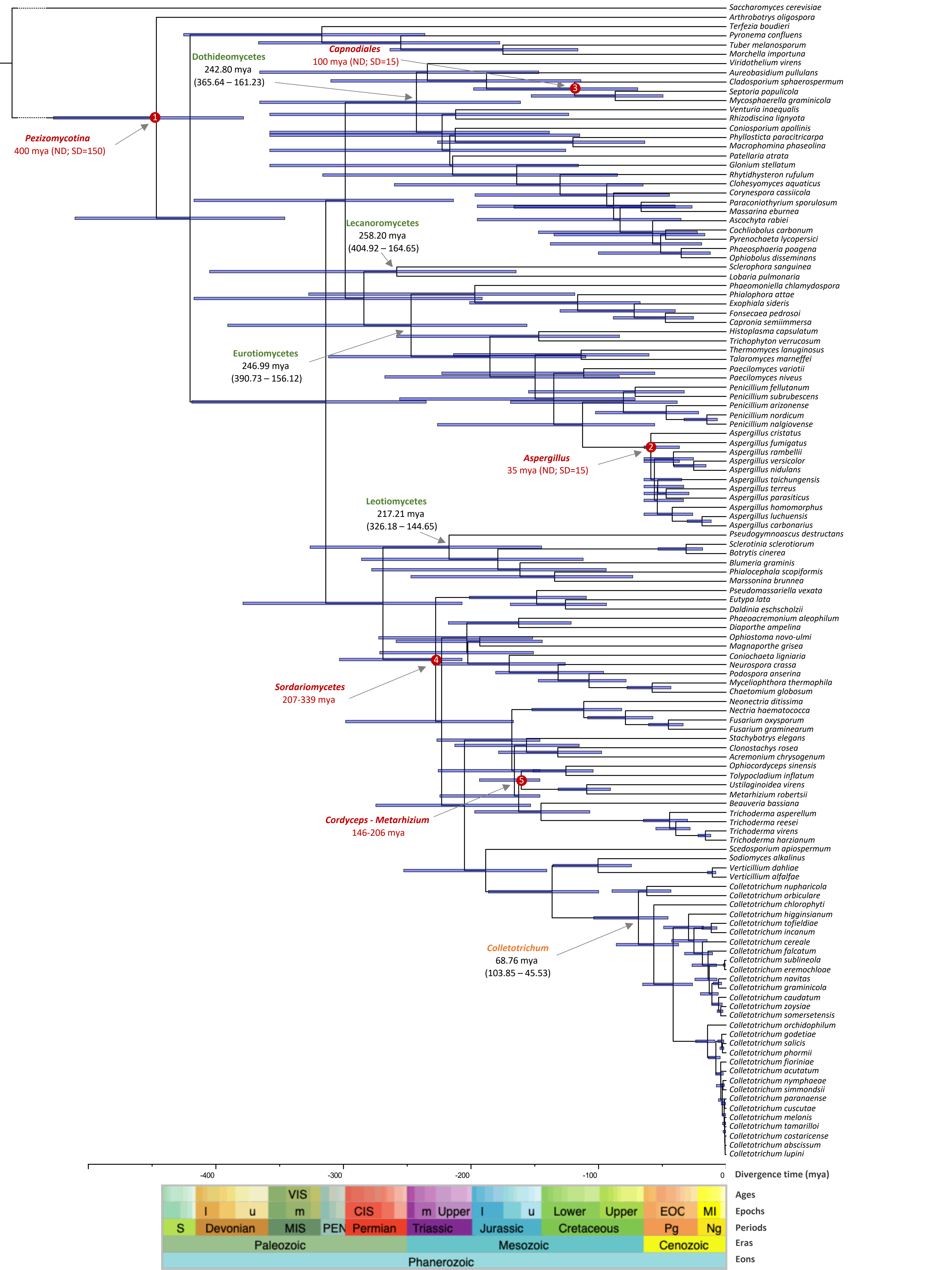

Saccharomyces cerevisiae
Arthrobotrys oligospora
Terfezia boudieri
Pyronema confluens
Tuber melanosporum
Morchella importuna
Viridothelium virens
Aureobasidium pullulans
Cladosporium sphaerospermum
Septoria populicola
Mycosphaerella graminicola
Venturia inaequalis
Rhizodiscina lignyota
Coniosporium apollinis
Phyllosticta paracitricarpa
Macrophomina phaseolina
Patellaria atrata
Glonium stellatum
Rhytidhysteron rufulum
Clohesyomyces aquaticus
Corynespora cassiicola
Paraconiothyrium sporulosum
Massarina eburnea
Ascochyta rabiei
Cochliobolus carbonum
Pyrenochaeta lycopersici
Phaeosphaeria poagena
Ophiobolus disseminans
Sclerophora sanguinea
Lobaria pulmonaria
Phaeomoniella chlamydospora
Phialophora attae
Exophiala sideris
Fonsecaea pedrosoi
Capronia semiimmersa
Histoplasma capsulatum
Trichophyton verrucosum
Thermomyces lanuginosus
Talaromyces marneffei
Paecilomyces variotii
Paecilomyces niveus
Penicillium fellutanum
Penicillium subrubescens
Penicillium arizonense
Penicillium nordicum
Penicillium nalgiovense
Aspergillus cristatus
Aspergillus fumigatus
Aspergillus rambellii
Aspergillus versicolor
Aspergillus nidulans
Aspergillus taichungensis
Aspergillus terreus
Aspergillus parasiticus
Aspergillus homomorphus
Aspergillus luchuensis
Aspergillus carbonarius
Pseudogymnoascus destructans
Sclerotinia sclerotiorum
Botrytis cinerea
Blumeria graminis
Phialocephala scopiformis
Marssonina brunnea
Pseudomassariella vexata
Eutypa lata
Daldinia eschscholzii
Phaeoacremonium aleophilum
Diaporthe ampelina
Ophiostoma novo-ulmi
Magnaporthe grisea
Coniochaeta ligniaria
Neurospora crassa
Podospora anserina
Myceliophthora thermophila
Chaetomium globosum
Neonectria ditissima
Nectria haematococca
Fusarium oxysporum
Fusarium graminearum
Stachybotrys elegans
Clonostachys rosea
Acremonium chrysogenum
Ophiocordyceps sinensis
Tolypocladium inflatum
Ustilaginoidea virens
Metarhizium robertsii
Beauveria bassiana
Trichoderma asperellum
Trichoderma reesei
Trichoderma virens
Trichoderma harzianum
Scedosporium apiospermum
Sodiomyces alkalinus
Verticillium dahliae
Verticillium alfalfae
Colletotrichum nupharicola
Colletotrichum orbiculare
Colletotrichum chlorophyti
Colletotrichum higginsianum
Colletotrichum tofieldiae
Colletotrichum incanum
Colletotrichum cereale
Colletotrichum falcatum
Colletotrichum sublineola
Colletotrichum eremochloae
Colletotrichum navitas
Colletotrichum graminicola
Colletotrichum caudatum
Colletotrichum zoysiae
Colletotrichum somersetensis
Colletotrichum orchidophilum
Colletotrichum godetiae
Colletotrichum salicis
Colletotrichum phormii
Colletotrichum fioriniae
Colletotrichum acutatum
Colletotrichum nymphaeae
Colletotrichum simmondsii
Colletotrichum paranaense
Colletotrichum cuscutae
Colletotrichum melonis
Colletotrichum tamarilloi
Colletotrichum costaricense
Colletotrichum abscissum
Colletotrichum lupini
Capnodiales
100 mya (ND; SD=15)
3
Dothideomycetes
242.80 mya
(365.64 – 161.23)
1
Pezizomycotina
400 mya (ND; SD=150)
Lecanoromycetes
258.20 mya
(404.92 – 164.65)
Eurotiomycetes
246.99 mya
(390.73 – 156.12)
2
Aspergillus
35 mya (ND; SD=15)
Leotiomycetes
217.21 mya
(326.18 – 144.65)
4
Sordariomycetes
207-339 mya
5
Cordyceps - Metarhizium
146-206 mya
Colletotrichum
68.76 mya
(103.85 – 45.53)
-400
-300
-200
-100
0
Divergence time (mya)
VIS
l
u
m
CIS
m
Upper
l
u
Lower
Upper
EOC
MI
S
Devonian
MIS
PEN
Permian
Triassic
Jurassic
Cretaceous
Pg
Ng
Paleozoic
Mesozoic
Cenozoic
Phanerozoic
Ages
Epochs
Periods
Eras
Eons

| Tree position | *Genus* | *species* | isolates | Database | Reference |
|---|---|---|---|---|---|
| 0 | *Saccharomyces* | *cerevisiae* | S288C | JGI | Goffeau A et al., 1996 |
| 1 | *Arthrobotrys* | *oligospora* | ATCC 24927 | JGI | Yang J et al., 2011 |
| 2 | *Terfezia* | *boudieri* | ATCC MYA-4762 | JGI | Murat C et al., 2018 |
| 3 | *Pyronema* | *confluens* | CBS 100304 | JGI | Traeger S et al., 2013 |
| 4 | *Tuber* | *melanosporum* | Mel28 v1.2 | JGI | Martin F et al., 2010 |
| 5 | *Morchella* | *importuna* | CCBAS932 | JGI | Murat C et al., 2018 |
| 6 | *Viridothelium* | *virens* | | JGI | Haridas et al., 2020 |
| 7 | *Aureobasidium* | *pullulans* | CBS 110374 | JGI | Gostincar C et al., 2014 |
| 8 | *Cladosporium* | *sphaerospermum* | UM 843 | JGI | Ng KP et al., 2012 |
| 9 | *Septoria* | *populicola* | | JGI | Ohm RA et al., 2012 |
| 10 | *Mycosphaerella* | *graminicola* | | JGI | Goodwin SB et al., 2011 |
| 11 | *Venturia* | *inaequalis* | | JGI | Deng CH et al., 2017 |
| 12 | *Rhizodiscina* | *lignyota* | CBS 133067 | JGI | Haridas et al., 2020 |
| 13 | *Coniosporium* | *apollinis* | CBS 100218 | JGI | Teixeira et al., 2017 |
| 14 | *Phyllosticta* | *paracitricarpa* | CBS 141357 | JGI | Guarnaccia et al., 2019 |
| 15 | *Macrophomina* | *phaseolina* | MS6 | JGI | Islam MS et al., 2012 |
| 16 | *Patellaria* | *atrata* | | JGI | Haridas et al., 2020 |
| 17 | *Glonium* | *stellatum* | CBS 207.34 | JGI | Peter M et al., 2016 |
| 18 | *Rhytidhysteron* | *rufulum* | | JGI | Ohm RA et al., 2012 |
| 19 | *Clohesyomyces* | *aquaticus* | | JGI | Mondo SJ et al., 2017 |
| 20 | *Corynespora* | *cassiicola* | CCP | JGI | Lopez D et al., 2018 |
| 21 | *Paraconiothyrium* | *sporulosum* | AP3s5-JAC2a | JGI | Zeiner CA et al., 2016 |
| 22 | *Massarina* | *eburnea* | CBS 473.64 | JGI | Haridas et al., 2020 |
| 23 | *Ascochyta* | *rabiei* | ArDII | JGI | Verma S et al., 2016 |
| 24 | *Cochliobolus* | *carbonum* | 26-R-13 | JGI | Condon BJ et al., 2013 |
| 25 | *Pyrenochaeta* | *lycopersici* | MPI-SDFR-AT-0127 | JGI | Mesny et al, 2021 |
| 26 | *Phaeosphaeria* | *poagena* | MPI-PUGE-AT-0046c | JGI | Mesny et al, 2021 |
| 27 | *Ophiobolus* | *disseminans* | CBS 113818 | JGI | Haridas et al., 2020 |
| 28 | *Sclerophora* | *sanguinea* | CBS 100924 | JGI | unpublished |
| 29 | *Lobaria* | *pulmonaria* | Scotland reference | JGI | unpublished |
| 30 | *Phaeomoniella* | *chlamydospora* | UCRPC4 | JGI | Morales-Cruz A et al., 2015 |
| 31 | *Phialophora* | *attae* | CBS 131958 | JGI | Moreno LF et al., 2015 |
| 32 | *Exophiala* | *sideris* | CBS 121828 | JGI | Teixeira MM et al., 2017 |
| 33 | *Fonsecaea* | *pedrosoi* | CBS 271.37 | JGI | Teixeira MM et al., 2017 |
| 34 | *Capronia* | *semiimmersa* | CBS 27337 | JGI | Teixeira MM et al., 2017 |
| 35 | *Histoplasma* | *capsulatum* | NAm1 | JGI | Sharpton TJ et al., 2009 |
| 36 | *Trichophyton* | *verrucosum* | HKI 517 | JGI | Burmester A et al., 2011 |
| 37 | *Thermomyces* | *lanuginosus* | SSBP | JGI | McHunu NP et al., 2013 |
| 38 | *Talaromyces* | *marneffei* | ATCC 18224 | JGI | Nierman et al., 2015 |
| 39 | *Paecilomyces* | *variotii* | CBS 101075 | JGI | Urquhart AS et al., 2018 |
| 40 | *Paecilomyces* | *niveus* | CO7 | JGI | Biango-Daniels et al., 2018 |
| 41 | *Penicillium* | *fellutanum* | ATCC 48694 | JGI | unpublished |
| 42 | *Penicillium* | *subrubescens* | CBS 132785 | JGI | Peng et al., 2017 |
| 43 | *Penicillium* | *arizonense* | CBS 141311 | JGI | Grijseels et al., 2016 |
| 44 | *Penicillium* | *nordicum* | DAOMC 185683 | JGI | Wingfield BD et al., 2015 |
| 45 | *Penicillium* | *nalgiovense* | FM193 | JGI | Nielsen JC et al., 2017 |
| 46 | *Aspergillus* | *cristatus* | GZAAS20.1005 | JGI | Ge Y et al., 2016 |
| 47 | *Aspergillus* | *fumigatus* | A1163 | JGI | Fedorova ND et al., 2008 |
| 48 | *Aspergillus* | *rambellii* | SRRC1468 | JGI | Moore et al., 2015 |
| 49 | *Aspergillus* | *versicolor* | | JGI | de Vries RP et al., 2017 |
| 50 | *Aspergillus* | *nidulans* | | JGI | Arnaud MB et al., 2012 |

| | | | | | |
|---|---|---|---|---|---|
| 51 | *Aspergillus* | *taichungensis* | IBT 19404 | JGI | Kjærbølling et al., 2018 |
| 52 | *Aspergillus* | *terreus* | NIH 2624 | JGI | Arnaud MB et al., 2012 |
| 53 | *Aspergillus* | *parasiticus* | CBS 117618 | JGI | Kjærbølling et al., 2020 |
| 54 | *Aspergillus* | *homomorphus* | CBS 101889 | JGI | Vesth TC et al., 2018 |
| 55 | *Aspergillus* | *luchuensis* | CBS 106.47 | JGI | de Vries RP et al., 2017 |
| 56 | *Aspergillus* | *carbonarius* | ITEM 5010 | JGI | de Vries RP et al., 2017 |
| 57 | *Pseudogymnoascus* | *destructans* | 20631-21 | JGI | Drees KP et al., 2016 |
| 58 | *Sclerotinia* | *sclerotiorum* | | JGI | Amselem J et al., 2011 |
| 59 | *Botrytis* | *cinerea* | | JGI | Staats M et al., 2012 |
| 60 | *Blumeria* | *graminis* | 96224 | JGI | Wicker T et al., 2013 |
| 61 | *Phialocephala* | *scopiformis* | 5WS22E1 | JGI | Walker AK et al., 2016 |
| 62 | *Marssonina* | *brunnea* | MB_m1 | JGI | Zhu S et al., 2012 |
| 63 | *Pseudomassariella* | *vexata* | CBS 129021 | JGI | Mondo SJ et al., 2017 |
| 64 | *Eutypa* | *lata* | UCREL1 | JGI | Blanco-Ulate B et al., 2013 |
| 65 | *Daldinia* | *eschscholzii* | EC12 | JGI | Wu W et al., 2017 |
| 66 | *Phaeoacremonium* | *aleophilum* | UCRPA7 | JGI | Blanco-Ulate B et al., 2013 |
| 67 | *Diaporthe* | *ampelina* | UCDDA912 | JGI | Morales-Cruz A et al., 2015 |
| 68 | *Ophiostoma* | *novo-ulmi* | H327 | JGI | Forgetta V et al., 2013 |
| 69 | *Magnaporthe* | *grisea* | | JGI | Dean RA et al., 2005 |
| 70 | *Coniochaeta* | *ligniaria* | NRRL 30616 | JGI | Jiménez et al., 2017 |
| 71 | *Neurospora* | *crassa* | OR74A | JGI | Galagan JE et al., 2003 |
| 72 | *Podospora* | *anserina* | S mat+ | JGI | Espagne E et al., 2008 |
| 73 | *Myceliophthora* | *thermophila* | | JGI | Berka RM et al., 2011 |
| 74 | *Chaetomium* | *globosum* | MPI-SDFR-AT-0079 | JGI | Mesny et al., 2021 |
| 75 | *Neonectria* | *ditissima* | R09/05 | JGI | Gomez-Cortecero A et al., 2015 |
| 76 | *Nectria* | *haematococca* | | JGI | Coleman JJ et al., 2009 |
| 77 | *Fusarium* | *oxysporum* | 4287 | JGI | Ma LJ et al., 2010 |
| 78 | *Fusarium* | *graminearum* | | JGI | Cuomo CA et al., 2007 |
| 79 | *Stachybotrys* | *elegans* | MPI-CAGE-CH-0235 | JGI | Mesny et al., 2021 |
| 80 | *Clonostachys* | *rosea* | IK726 | JGI | Karlsson et al., 2015 |
| 81 | *Acremonium* | *chrysogenum* | ATCC 11550 | JGI | Terfehr D et al., 2014 |
| 82 | *Ophiocordyceps* | *sinensis* | IOZ07 | NCBI | Shu et al., 2020 |
| 83 | *Tolypocladium* | *inflatum* | NRRL 8044 | JGI | Bushley KE et al., 2013 |
| 84 | *Ustilaginoidea* | *virens* | | JGI | Kumagai T et al., 2016 |
| 85 | *Metarhizium* | *robertsii* | ARSEF 23 | NCBI | Hu et al., 2014 |
| 86 | *Beauveria* | *bassiana* | ARSEF 2860 | JGI | Xiao G et al., 2012 |
| 87 | *Trichoderma* | *asperellum* | CBS 433.97 | JGI | Druzhinina IS et al., 2018 |
| 88 | *Trichoderma* | *reesei* | QM6a | JGI | Li WC et al., 2017 |
| 89 | *Trichoderma* | *virens* | Gv29-8 | JGI | Kubicek CP et al., 2011 |
| 90 | *Trichoderma* | *harzianum* | TR274 | JGI | Kubicek CP et al., 2019 |
| 91 | *Scedosporium* | *apiospermum* | IHEM 14462 | JGI | Vandeputte et al., 2014 |
| 92 | *Sodiomyces* | *alkalinus* | | JGI | Grum-Grzhimaylo AA et al., 2018 |
| 93 | *Verticillium* | *dahliae* | VdLs.17 | JGI | Klosterman SJ et al., 2011 |
| 94 | *Verticillium* | *alfalfae* | VaMs.102 | JGI | Klosterman SJ et al., 2011 |
| 95 | *Colletotrichum* | *nupharicola* | 23 | JGI | unpublished |
| 96 | *Colletotrichum* | *orbiculare* | MAFF 240422 | JGI | Gan P et al., 2013 |
| 97 | *Colletotrichum* | *chlorophyti* | NTL11 | JGI | Gan P et al., 2017 |
| 98 | *Colletotrichum* | *higginsianum* | IMI 349063 | JGI | Zampounis A et al., 2016 |
| 99 | *Colletotrichum* | *tofieldiae* | 861 | JGI | Hackard et al., 2016 |
| 100 | *Colletotrichum* | *incanum* | MAFF 238712 | JGI | Gan P et al., 2016 |
| **101** | ***Colletotrichum*** | ***cereale*** | **CBS 129662** | **JGI** | **This work** |
| **102** | ***Colletotrichum*** | ***falcatum*** | **MAFF 306170** | **JGI** | **This work** |

| | | | | | |
|---|---|---|---|---|---|
| **103** | ***Colletotrichum*** | ***sublineola*** | **CBS 131301** | **JGI** | **This work** |
| **104** | ***Colletotrichum*** | ***eremochloae*** | **CBS 129661** | **JGI** | **This work** |
| **105** | ***Colletotrichum*** | ***navitas*** | **CBS 125086** | **JGI** | **This work** |
| 106 | *Colletotrichum* | *graminicola* | M1.001 | JGI | O'Connell RJ et al., 2012 |
| **107** | ***Colletotrichum*** | ***caudatum*** | **CBS 131602** | **JGI** | **This work** |
| **108** | ***Colletotrichum*** | ***zoysiae*** | **MAFF 235873** | **JGI** | **This work** |
| **109** | ***Colletotrichum*** | ***somersetensis*** | **CBS 131599** | **JGI** | **This work** |
| 110 | *Colletotrichum* | *orchidophilum* | IMI 309357 | JGI | Baroncelli R et al., 2018 |
| **111** | ***Colletotrichum*** | ***godetiae*** | **CBS 193.32** | **JGI** | **This work** |
| 112 | *Colletotrichum* | *salicis* | CBS 607.94 | JGI | Baroncelli R et al., 2016 |
| **113** | ***Colletotrichum*** | ***phormii*** | **CBS 102054** | **JGI** | **This work** |
| 114 | *Colletotrichum* | *fioriniae* | IMI 504882 | JGI | Baroncelli R et al., 2014 |
| **115** | ***Colletotrichum*** | ***acutatum*** | **CBS 112980** | **JGI** | **This work** |
| 116 | *Colletotrichum* | *nymphaeae* | IMI 504889 | JGI | Baroncelli R et al., 2016 |
| 117 | *Colletotrichum* | *simmondsii* | CBS 122122 | JGI | Baroncelli R et al., 2016 |
| **118** | ***Colletotrichum*** | ***paranaense*** | **IMI 384185** | **JGI** | **This work** |
| 119 | *Colletotrichum* | *cuscutae* | IMI 304802 | JGI | Baroncelli et al., 2021 |
| **120** | ***Colletotrichum*** | ***melonis*** | **CBS 134730** | **JGI** | **This work** |
| **121** | ***Colletotrichum*** | ***tamarilloi*** | **CBS 129955** | **JGI** | **This work** |
| **122** | ***Colletotrichum*** | ***costaricense*** | **IMI 309622** | **JGI** | **This work** |
| **123** | ***Colletotrichum*** | ***abscissum*** | **IMI 504890** | **JGI** | **This work** |
| **124** | ***Colletotrichum*** | ***lupini*** | **CBS 109225** | **JGI** | **This work** |
