## Supplementary material for "Genome evolution and transcriptome plasticity associated with adaptation to monocot and eudicot plants in *Colletotrichum* fungi": Suppl11_Evo_genefamilies.pdf

### PL-6 family (IPR039513)

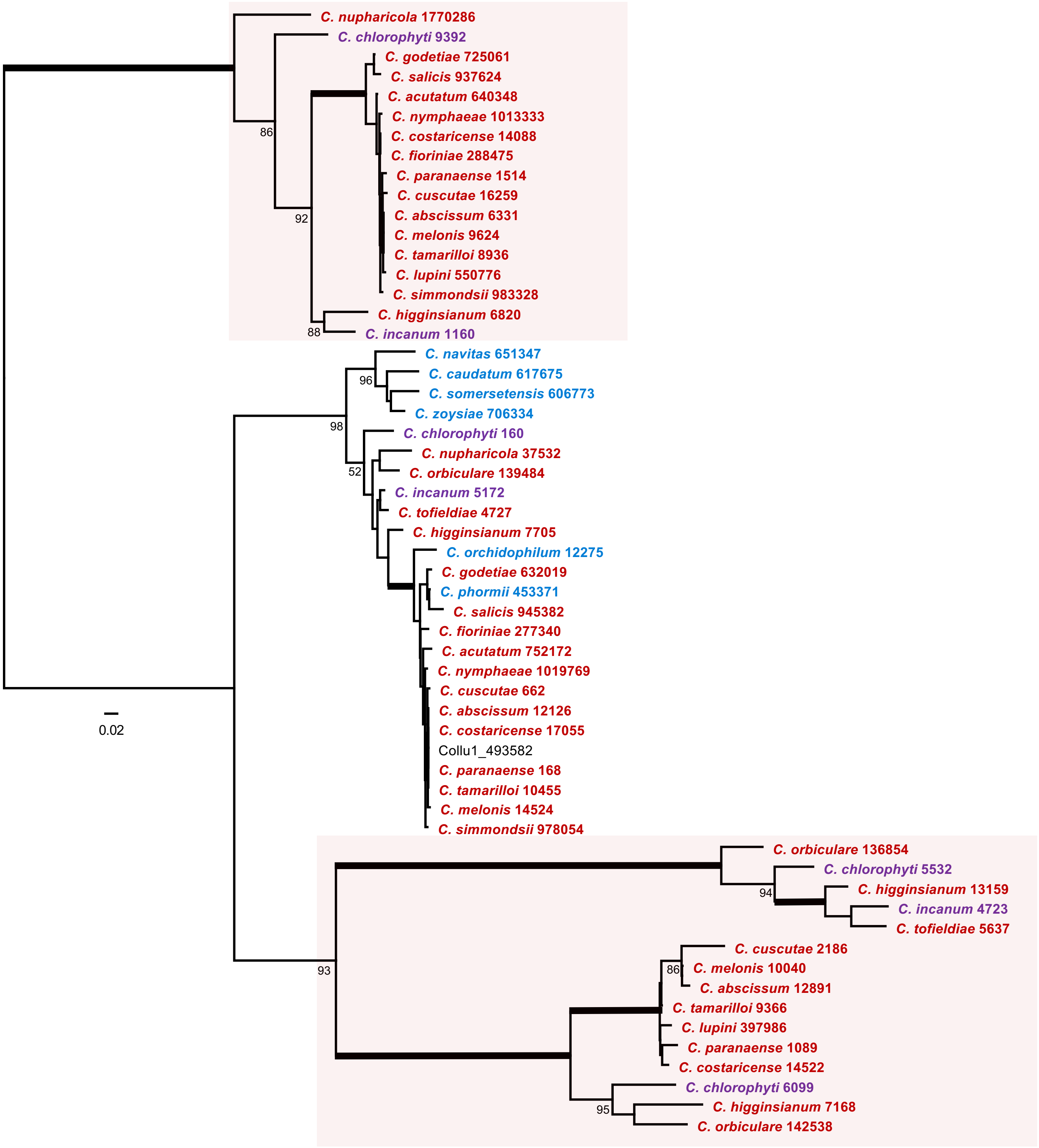

### Transcription initiation factor IID, subunit 13 (IPR003195)

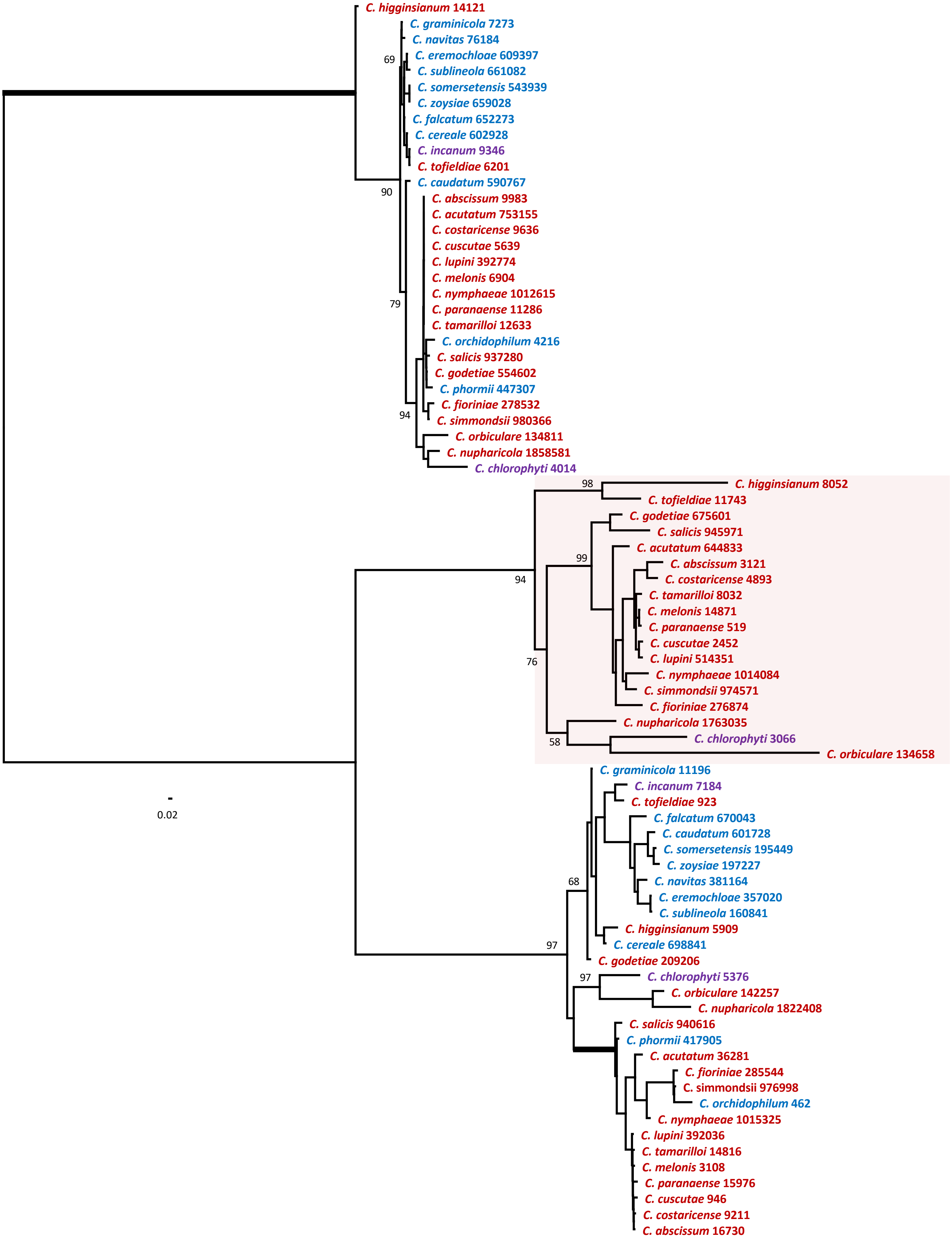

### Aconitase, mitochondrial-like (IPR006248)

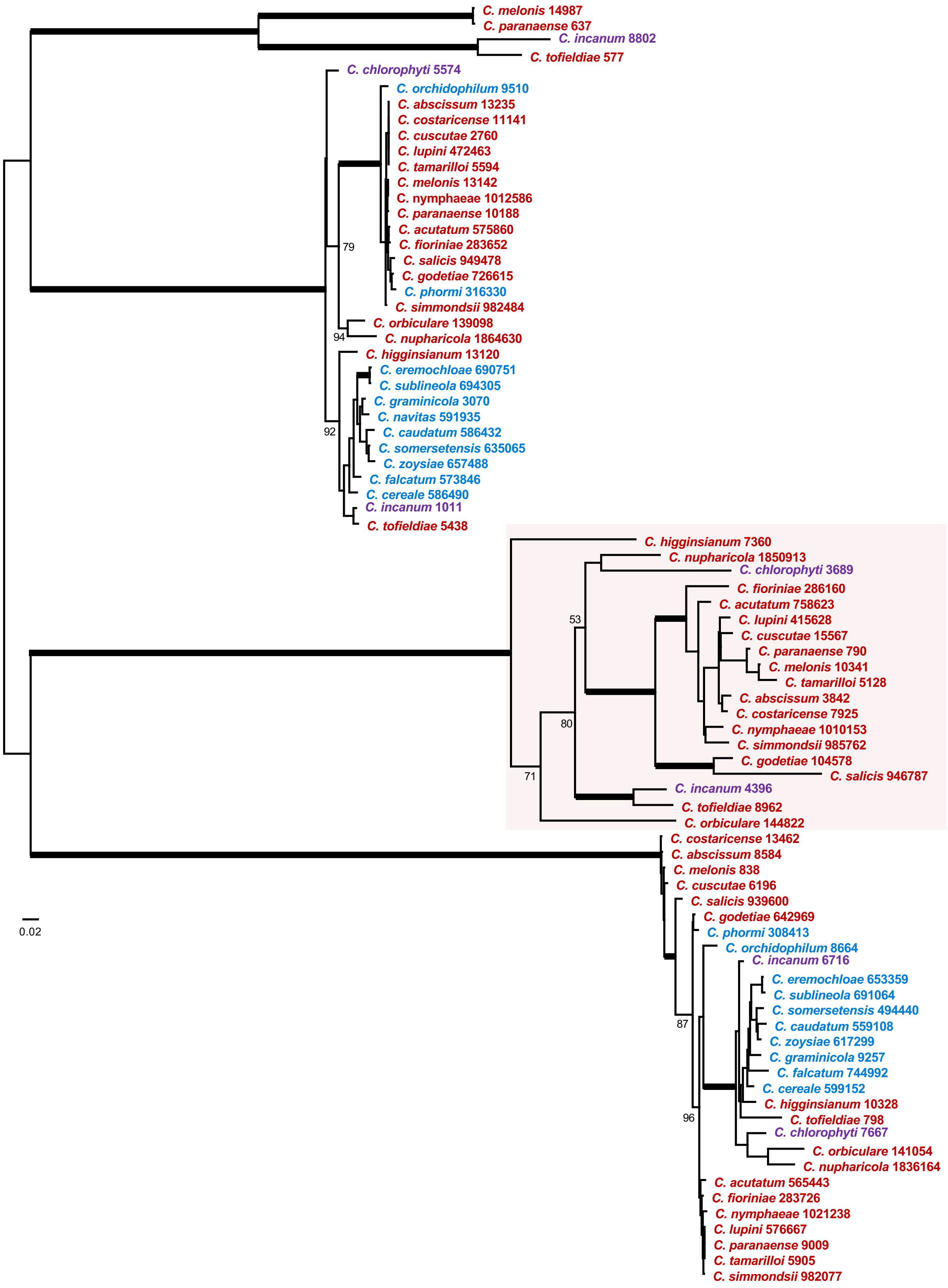

PoSI-like peptidase domain (IPR034187)

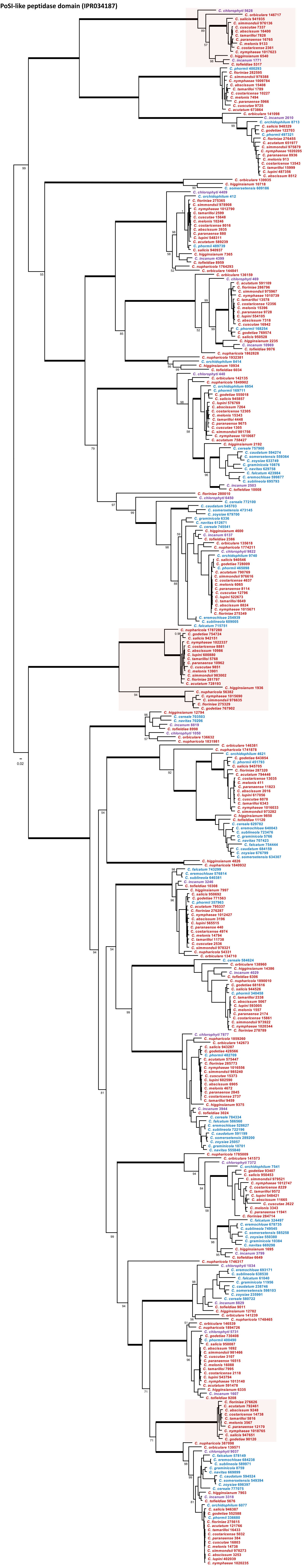
