## Supplementary figures and images for "Genome evolution and transcriptome plasticity associated with adaptation to monocot and eudicot plants in *Colletotrichum* fungi"

### Suppl12_replicates_heatmaps.pdf

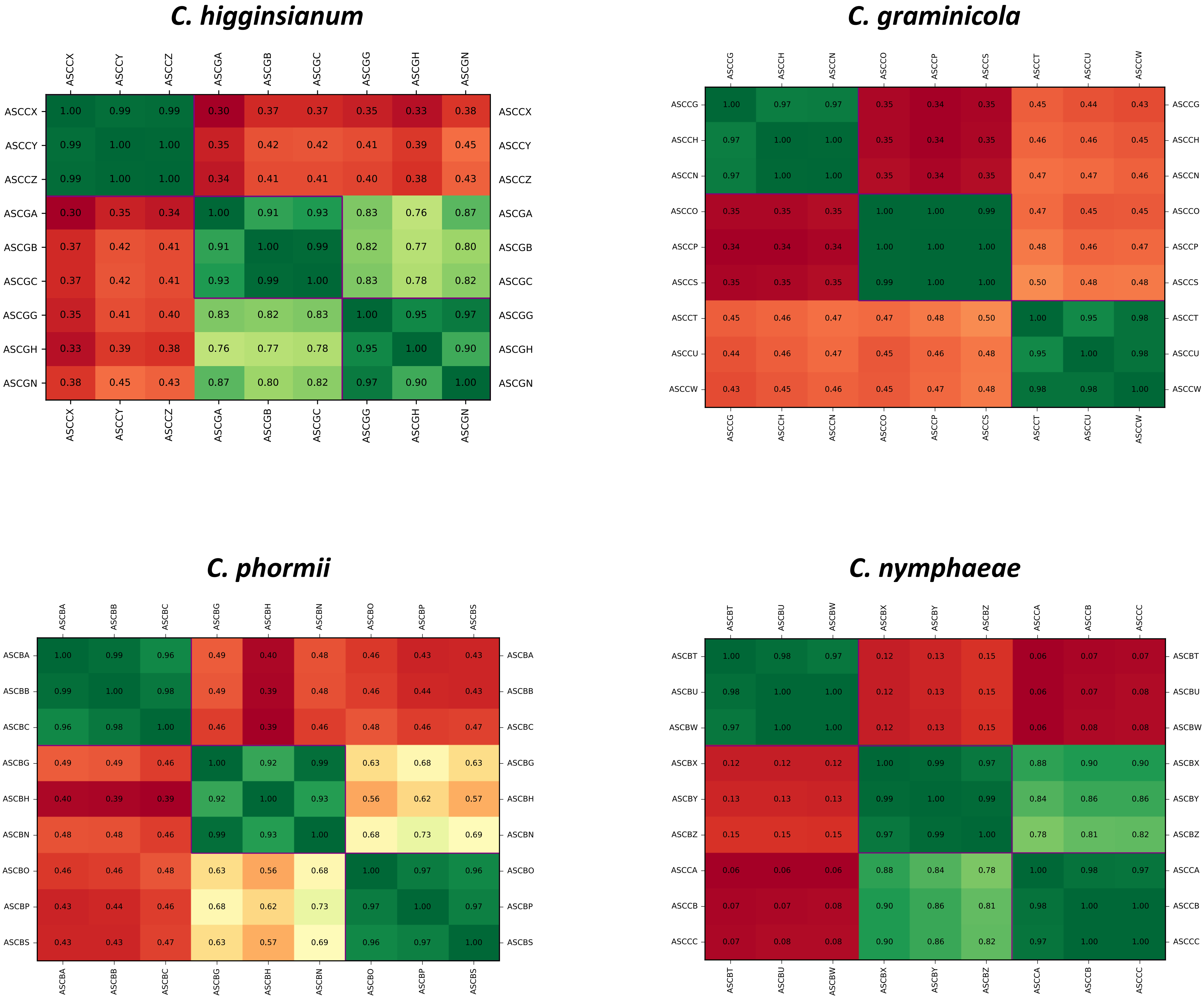

### Suppl13_Vulcano_plots.pdf

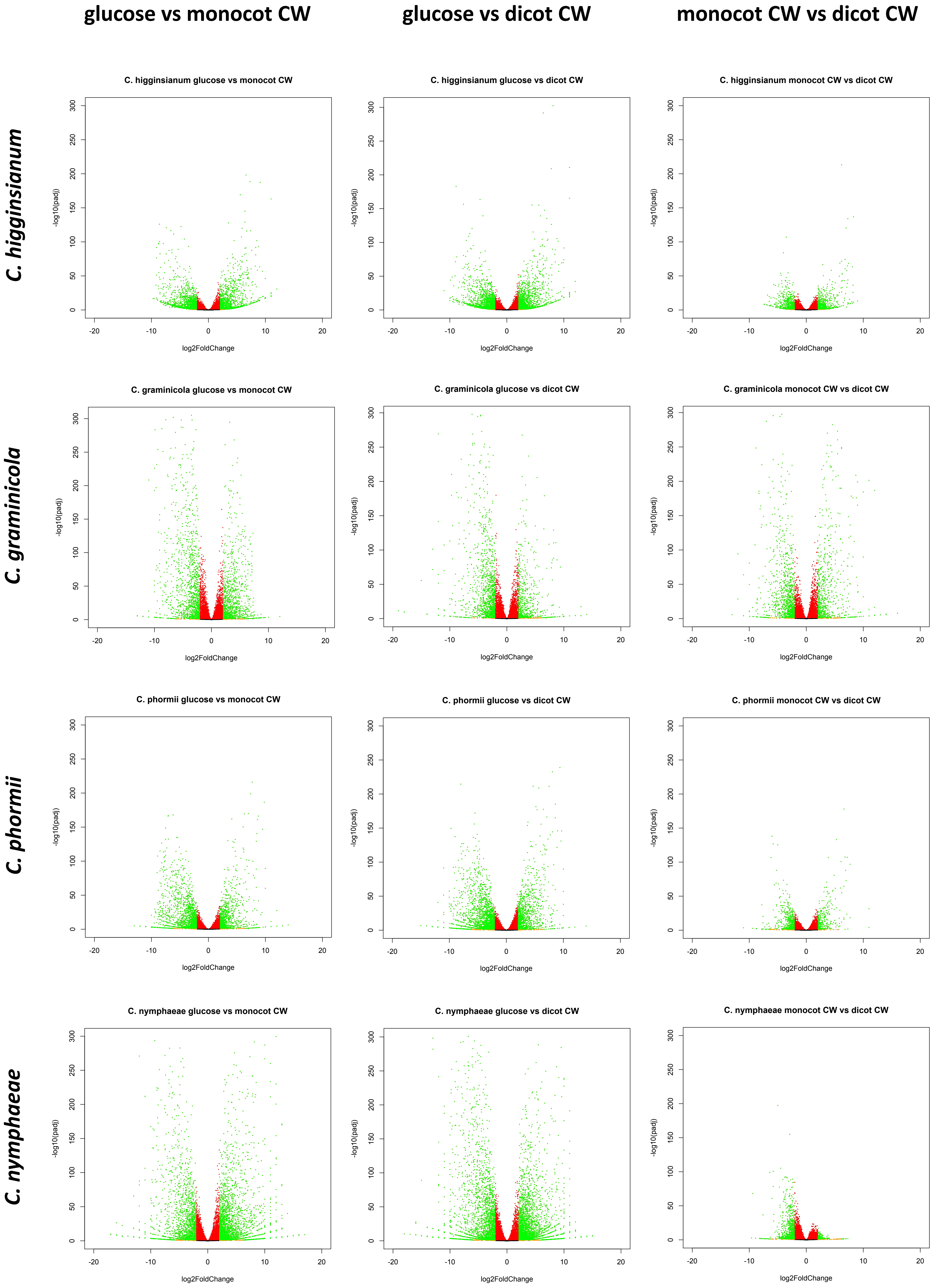
